# Prophage-encoded methyltransferase drives adaptation of community-acquired methicillin-resistant *Staphylococcus aureus*

**DOI:** 10.1101/2024.04.17.589803

**Authors:** Robert J. Ulrich, Magdalena Podkowik, Rebecca Tierce, Irnov Irnov, Gregory Putzel, Nora Samhadaneh, Keenan A. Lacey, Daiane Boff, Sabrina M. Morales, Sohei Makita, Theodora K. Karagounis, Erin E. Zwack, Chunyi Zhou, Randie Kim, Karl Drlica, Alejandro Pironti, Harm van Bakel, Victor J. Torres, Bo Shopsin

## Abstract

We recently described the evolution of a community-acquired methicillin-resistant *Staphylococcus aureus* (CA-MRSA) USA300 variant responsible for an outbreak of skin and soft tissue infections. Acquisition of a mosaic version of the Φ11 prophage (mΦ11) that increases skin abscess size was an early step in CA-MRSA adaptation that primed the successful spread of the clone. The present report shows how prophage mΦ11 exerts its effect on virulence for skin infection without encoding a known toxin or fitness genes. Abscess size and skin inflammation were associated with DNA methylase activity of an mΦ11-encoded adenine methyltransferase (designated *pamA*). *pamA* increased expression of fibronectin-binding protein A (*fnbA*; FnBPA), and inactivation of *fnbA* eliminated the effect of *pamA* on abscess virulence without affecting strains lacking *pamA*. Thus, *fnbA* is a *pamA*-specific virulence factor. Mechanistically, *pamA* was shown to promote biofilm formation in vivo in skin abscesses, a phenotype linked to FnBPA’s role in biofilm formation. Collectively, these data reveal a novel mechanism—epigenetic regulation of staphylococcal gene expression—by which phage can regulate virulence to drive adaptive leaps by *S. aureus*.

**Graphical Abstract:** 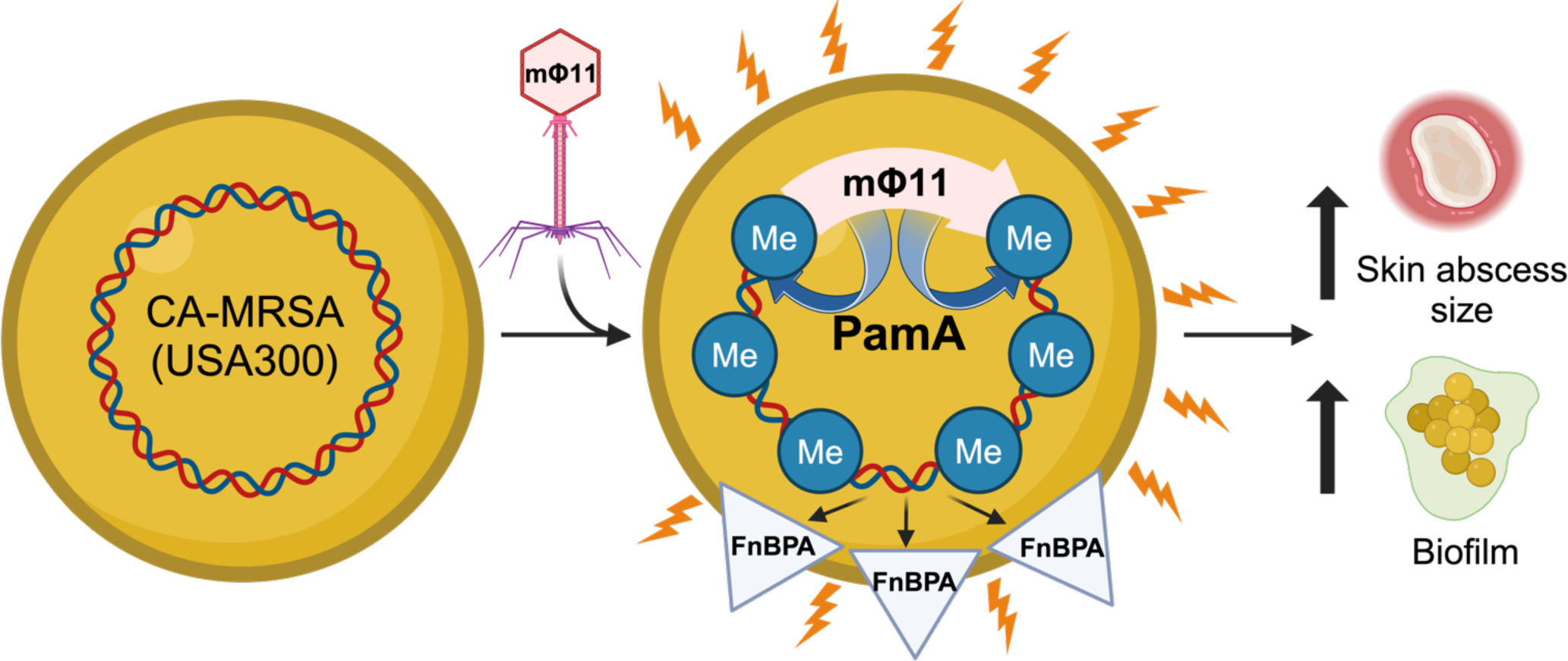

## Introduction

Community-acquired methicillin-resistant *Staphylococcus aureus* (CA-MRSA) lineage USA300 is the major cause of skin and soft tissue infection in the United States, and clonal variants that cause outbreaks have become public health emergencies (1-3). The spectrum of adaptive changes that arise during the course of CA-MRSA dissemination is likely to identify genetic pathways critical for bacterial pathogenesis in vivo (4). We recently described a genotypic cluster of CA-MRSA-USA300 that emerged in Brooklyn, NY (USA300-BKV) that was uniquely positioned to offer insight into properties of emerging CA-MRSA strains (5). The persistence of the Brooklyn disease cluster enabled us to use phylogenetic analysis and experimental assays to identify a unique prophage that promoted large skin abscesses. That pathogen advantage primed USA300-BKV for successful spread, thereby facilitating the subsequent emergence of resistance to topical antimicrobials (5). Until now, it was unknown how the Brooklyn cluster-associated prophage, which is a mosaic variant of the well-known *S. aureus* generalized transducing phage Φ11 (referred to as mΦ11), enhanced virulence during skin infection.

*S. aureus* strains often carry multiple prophages (6-8), which are primary drivers of *S. aureus* evolution, diversity, and virulence (9-12). To date, studies of prophage-mediated virulence in *S. aureus* have primarily focused on prophage-encoded toxin and fitness genes. One example is the prophage-encoded Panton-Valentine leucocidin which is associated with skin abscesses across all *S. aureus* lineages, including USA300 (13-16). In contrast, the Brooklyn cluster-associated mΦ11 is similar to most prophages in that it lacks a known virulence factor, nor does its insertion disrupt a chromosomal virulence gene (5). Thus, new pathogenic mechanisms are expected to derive from genetic and phenotypic analysis that define mΦ11 components driving enhanced skin abscess virulence.

In the present work, we report that the methylase activity of an mΦ11-encoded DNA methyltransferase, which has been named *pamA* for phage adenine methyltransferase A, is necessary and sufficient for the enhanced skin abscess phenotype observed with the USA300-BKV clone. Moreover, we found that *pamA* rewires the bacterial transcriptional program, resulting in a marked increase in the expression and production of fibronectin-binding protein A (*fnbA*; FnBPA), as determined by RNA sequencing and global proteomics. *fnbA* was necessary for the *pamA*-associated skin abscess phenotype, which was in turn associated with increased production of biofilms. Collectively, the data demonstrate how phage can modify DNA to enhance USA300 virulence by altering the expression of core genome-encoded virulence factors, thereby increasing the fitness of epidemic clones and driving leaps in adaptation.

## Results

### Genetic deletions localize the mΦ11 gene(s) responsible for increased skin infection virulence

Prophage mΦ11 promotes USA300-mediated tissue damage by increasing skin abscess size during murine infection **(Figure S1A)**, as previously reported (5). Often, hypervirulent strains of *S. aureus* will exhibit differences in growth rates (17), secreted protein production (18), and/or transcriptional profiles (19-21). However, strain USA300 LAC* harboring mΦ11 did not show significantly altered in vitro growth kinetics, exoprotein production, hemolysis patterns, or transcript levels of non-mΦ11 genes compared to the parental USA300 LAC*, as determined by RNA sequencing analysis **(Figures S1B-E)**. Therefore, results of the in vitro analyses did not correlate with the increased virulence demonstrated by mΦ11-containing strains during skin infection. These data suggest that an in vivo signal(s) is required for mΦ11-associated virulence (22). Thus, in vivo models of infection are required to identify mechanisms underlying mΦ11-mediated virulence **(Figure 1A)**.

**Figure 1.**
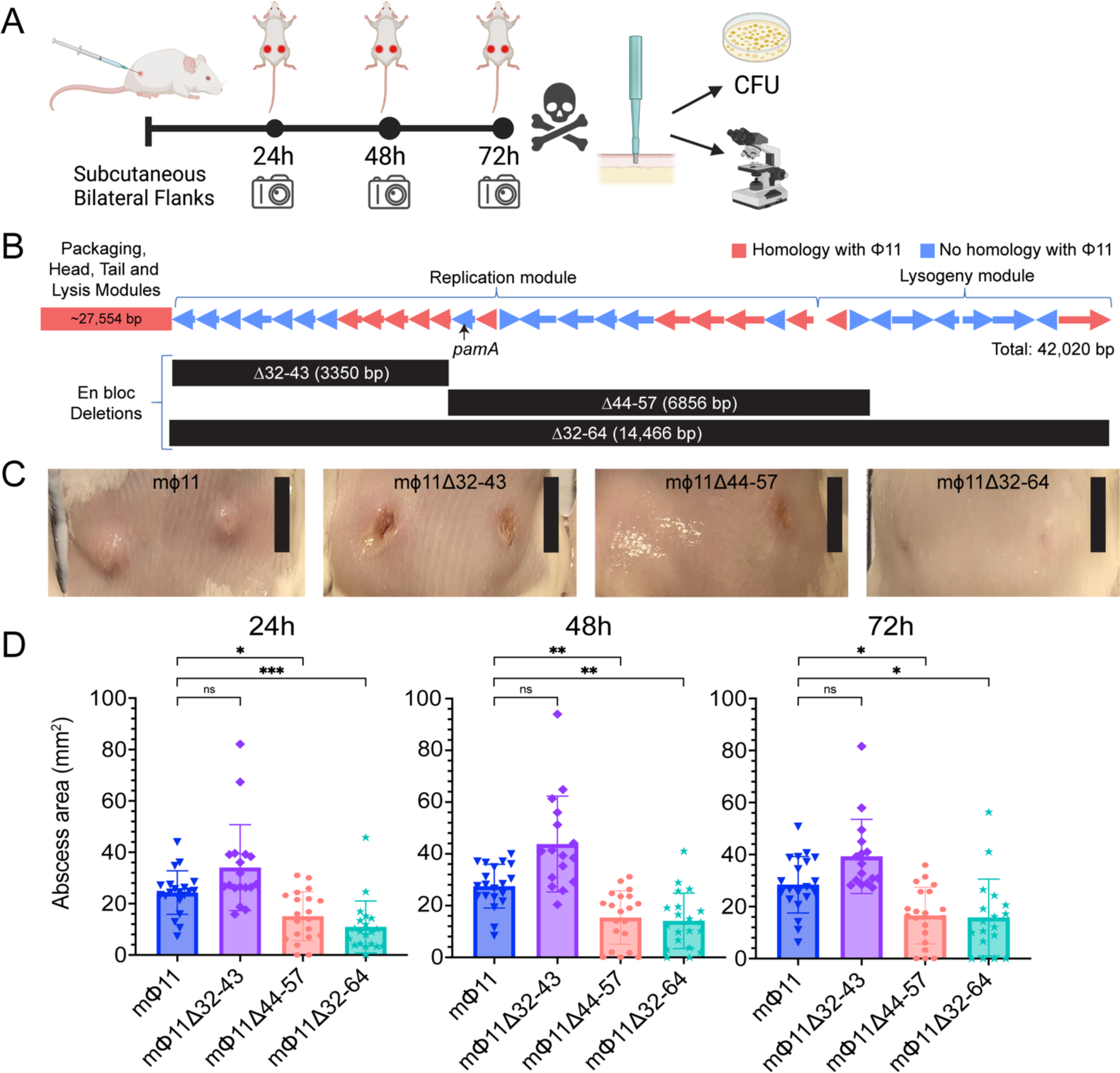
Effect of en block deletions on the mΦ11-mediated skin abscess phenotype. **(A)** Schematic of skin infection workflow. CFU, colony forming units. Created with BioRender.com. **(B)** Map of mΦ11 in strain USA300-BKV, adapted with permission from (5), with en bloc deletion locations. Arrows indicate predicted ORFs and the direction of the transcription of genes within the unique mΦ11 modules. Homologous (red) and non-homologous (blue) ORFs are shown, as compared to prototypical Φ11. Black arrow indicates *pamA*. Black bars beneath the gene map correspond to the gene blocks deleted from the indicated strain. **(C)** Representative images of skin abscesses 72 h after subcutaneous infection with the indicated strains. Scale bar (black) is 1cm. **(D)** Skin abscess infections with en bloc deletion mutants. Skin abscess area of LAC* lysogens containing mΦ11 (blue, N=20, strain BS989), mΦ11Δ32-43 (purple, N=16-18, strain RU47), mΦ11Δ44-57 (salmon, N=20, strain RU108) and mΦ11Δ32-64 (cyan, N=18-20, strain RU42) at 24, 48, and 72 hours after subcutaneous infection with ∼10^7^ CFU of bacteria. Data are pooled from two independent experiments and represent mean ± SD. Statistical significance was determined with the Kruskall-Wallis test and Dunn’s multiple comparisons test, \**P*≤.05, ** *P*≤.01, \*\*\**P*≤.001.

Annotation of the mosaic portion of mΦ11 failed to identify known virulence factors (5). Consequently, we constructed three en bloc deletions within the mosaic region of mΦ11 to identify candidate gene(s) **(Figure 1B)**. The deletions were confirmed using whole genome sequencing **(Figure S2).** As expected, deletion of the entire mosaic region containing genes in the replication and lysogeny modules (Δ32-64) eliminated the mΦ11 skin abscess phenotype in mice **(Figures 1C-D).** Although deletion of an upstream fragment (Δ32-43) had no impact on abscess size, deletion of the center gene block (Δ44-57) eliminated the skin abscess phenotype **(Figures 1C-D)**. These data localized skin abscess candidates to 14 genes (44-57) in mΦ11 for further analysis. Of the 14 genes, eight gene sequences were unrelated to prototypical Φ11 or other known prophages and therefore were considered promising candidates for further analysis **(Table S1)**.

### An mΦ11-encoded adenine methyltransferase (pamA) is responsible for increased skin infection virulence

Examination of the eight potential virulence genes identified a methyltransferase that was absent in wild-type Φ11 (**Table S1**). The mΦ11-encoded adenine methyltransferase (*pamA*) shares amino acid sequence homology with DNA adenine methyltransferases (*dam*) (5), so called orphan methyltransferases that are not paired with a cognate restriction endonuclease and therefore do not form an obvious restriction-modification system. DNA adenine methyltransferases act independently to regulate gene expression and bacterial replication (23-25). They have also been implicated in prophage-mediated pathogenicity of an outbreak strain of *E. coli* (26). Thus, *pamA* represented a promising candidate gene as a novel virulence factor.

To determine whether *pamA* is necessary for the enhanced skin abscess phenotype, we engineered an in-frame, unmarked *pamA* deletion in a USA300 LAC* mΦ11 lysogen (mΦ11Δ*pamA*). Sanger and whole-genome sequencing respectively confirmed the deletion and the absence of adventitious secondary mutations in mΦ11 Δ*pamA* **(Figure S3).** Infection of mice with mΦ11Δ*pamA* resulted in a nearly identical average skin abscess size compared to that of the control Φ11 lysogen **(Figure 2A)**, suggesting that *pamA* was necessary for increased virulence. Complementation, by integration of constitutively expressed *pamA* into the staphylococcal chromosome in single copy using the SaPI *att* site (27), confirmed that *pamA* is responsible for the skin abscess phenotype **(Figure 2C)**. We did not observe a difference in bacterial burden at 72 h post-infection **(Figures 2B, 2D)**, as previously reported for comparisons between mΦ11 and Φ11 lysogens (5). Collectively, these data demonstrate that mΦ11-encoded *pamA* is required for increased skin abscess size in mice.

**Figure 2.**
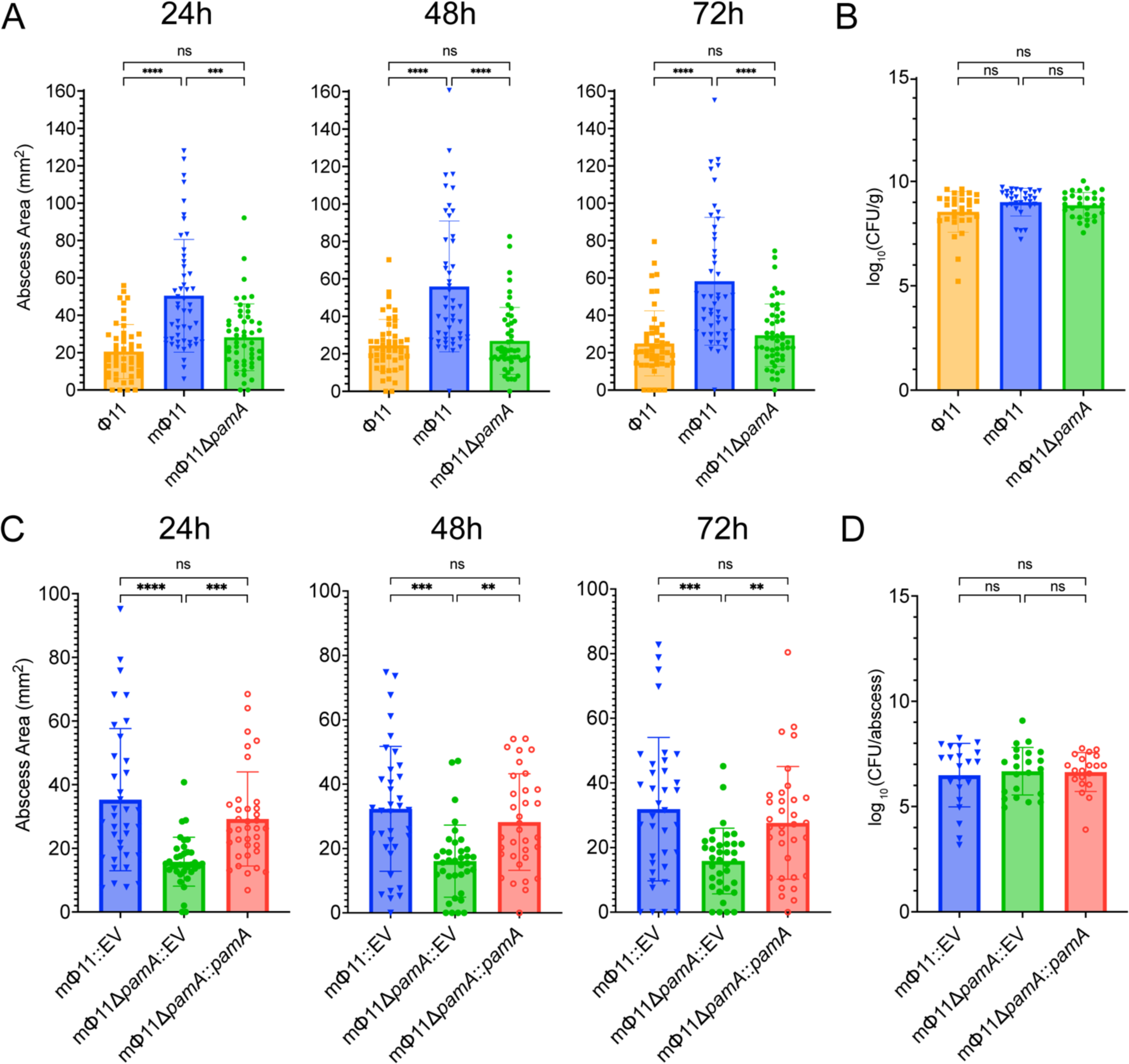
mΦ11 phage adenine methyltransferase (*pamA*) increases skin abscess size without affecting tissue bacterial burden. **(A)** Effect of *pamA* on skin abscess size. Abscess area of LAC* lysogens containing Φ11 (orange, N=50 abscesses, strain BS990), mΦ11 (blue, N=48-50 abscesses, strain BS989), and mΦ11Δ*pamA* (green, N=50 abscesses, strain RU39) at 24, 48, and 72 h after infection with ∼1.5x10^7^ CFU of bacteria per abscess. Results are pooled from four independent experiments. Data represent mean ± SD. Statistical significance was determined with the Kruskall-Wallis test and Dunn’s multiple comparisons test, \*\*\**P*≤.001, \*\*\*\**P*≤.0001. **(B)** *pamA* skin abscess phenotype and bacterial burden. Skin abscesses infections with Φ11 (orange, N=30 abscesses, strain BS990), mΦ11 (blue, N=30 abscesses, strain BS989), or mΦ11Δ*pamA* (green, N=30 abscesses, strain RU39) lysogens in LAC*. CFU were enumerated at 72 h. Data represent mean ± SD. Statistical significance was determined with the Kruskall-Wallis test and Dunn’s multiple comparisons test. **(C)** Effect of *pamA* complementation on abscess size. Abscess area of LAC* containing mΦ11::EV (blue, N=36 abscesses, strain RU138), mΦ11Δ*pamA*::EV (green, N=36 abscesses, strain RU128), mΦ11Δ*pamA*::*pamA* (red, N=34-36 abscesses, strain RU131) after infection with ∼1x10^7^ CFU of bacteria for the indicated times. EV, empty vector. Results are pooled from four independent experiments. Data represent mean ± SD. Statistical significance was determined with the Kruskall-Wallis test and Dunn’s multiple comparisons test, \*\*\**P*≤.001, \*\*\*\**P*≤.0001. **(D)** Bacterial burden in abscesses. Skin abscesses of LAC* containing mΦ11::EV (blue, N=22 abscesses, strain RU138), mΦ11Δ*pamA*::EV (green, N=22 abscesses, strain RU128), and mΦ11Δ*pamA*::*pamA* (red, N=20 abscesses, strain RU131) were harvested at 72 h and CFU enumerated. Of note, CFU/abscess is shown due to missing abscess weights during one of the replicate experiments. With the available weight adjusted data we found no significant differences between strains (data not shown). Data represent mean ± SD. Statistical significance was determined with the Kruskall-Wallis test and Dunn’s multiple comparisons test.

### pamA increases CA-MRSA skin abscess size irrespective of other mΦ11 genes

Next, we hypothesized that *pamA* expression would increase CA-MRSA virulence independent of other mΦ11 genes. Indeed, wild-type USA300 LAC* expressing *pamA* (LAC*::*pamA*) produced larger abscesses than an empty vector control strain (LAC*::EV) at all times post-infection **(Figure 3A)**. A maximal increase in abscess area of 91% was observed at 48 h post-infection. Therefore, *pamA* is sufficient to increase CA-MRSA skin virulence. As with mΦ11 lysogens, LAC*::*pamA* did not affect bacterial CFU recovered from skin abscesses **(Figure 3B)**, supporting the hypothesis that *pamA* increases abscess size by increasing tissue inflammation rather than bacterial burden. To test if this hypothesis is true, we compared skin abscess histology and murine cytokine production of LAC*::*pamA* and LAC*::EV at 72 h post-infection. Skin abscess inflammatory burden **(Figures 3C-D)** and proinflammatory cytokine/chemokine production **(Figure 3E)** were increased in LAC*::*pamA* skin abscesses compared to control LAC*:EV. Together, these data demonstrate that insertion of *pamA* into a USA300 LAC* background without the surrounding mΦ11 genes is sufficient to increase local tissue inflammation and, thereby, skin abscess size.

**Figure 3.**
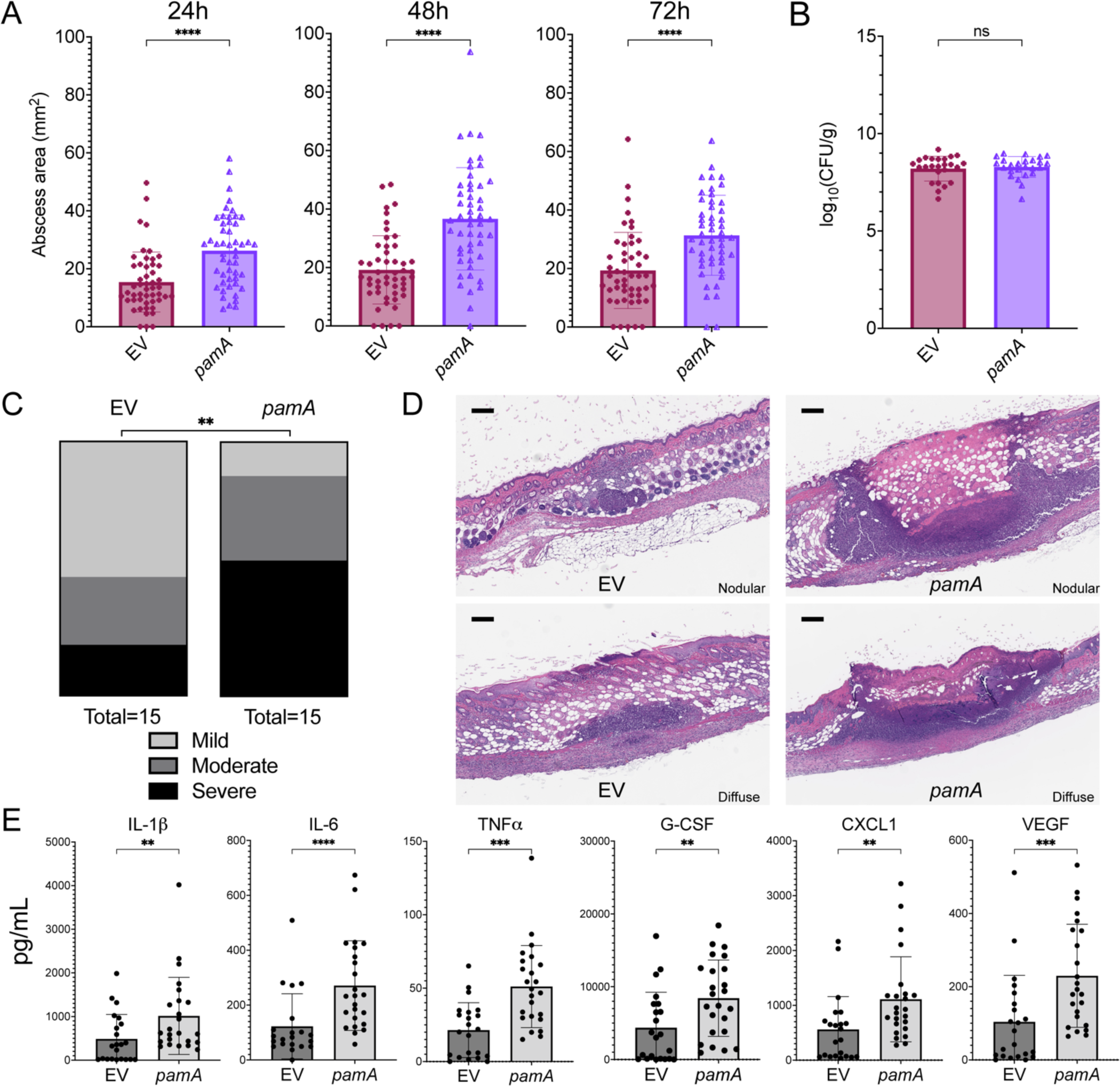
*pamA* increases skin abscess size and inflammation in the absence of mΦ11. **(A)** Effect of *pamA* on skin abscess size. Abscess area of LAC* with empty vector (EV) (maroon, N=50 abscesses, strain RU129) or constitutively expressed *pamA* (purple, N=50 abscesses, strain RU121) integrated into the chromosome in single copy after infection in with ∼1x10^7^ CFU of bacteria per abscess for the indicated times. Data are pooled from four independent experiments and represent mean ± SD. Statistical significance was determined with the Mann-Whitney test, \*\*\*\**P*≤.0001. **(B)** Effect of *pamA* on CFU recovered from skin abscesses. Skin abscesses (N=25 abscesses per strain) from two independent infections in panel A were harvested at 72h. Data represent mean ± SD of CFU recovered. Statistical significance was determined with the Mann-Whitney test. **(C)** Effect of *pamA* on skin inflammation. Biopsies of skin abscess (N=15 per strain, pooled from two independent experiments) from LAC* containing EV (strain RU129) *or pamA* (strain RU121) were stained with H&E and inflammatory burden graded by a blinded dermatopathologist. Statistical significance was determined with chi-square test (*P* = 0.0014). **(D)** Representative images of skin abscess biopsies from panel C. One representative image from each strain is presented according to dermatopathologist classification as nodular (above) or diffuse (below) architecture. **(E)** Effect of *pamA* on local proinflammatory and vascular proliferation cytokines. Biopsy of skin abscesses from three independent experiments of LAC* with EV control (N=22 abscesses, strain RU129) or *pamA* (N=24 abscesses, strain RU121) were homogenized and the indicated cytokine levels measured. Data represent mean ± SD. Statistical significance was determined with the Mann-Whitney test, ** *P*≤.01, \*\*\**P*≤.001, \*\*\*\**P*≤.0001.

### pamA-associated skin abscess virulence depends on methyltransferase activity

To determine whether *pamA* increases skin abscess virulence through methylase activity, we identified the conserved Dam active site (NPPY) in PamA **(Figure 4A)** and individually introduced several point mutations in residues previously reported to inactivate methyltransferase activity (28). To confirm that the PamA point mutants were inactive, we digested genomic DNA with the restriction endonuclease DpnI, an enzyme that digests at the methylated target of Dam (G<u>A</u>TC) (29). As expected, *pamA*-containing strains, but not those with point mutations in *pamA,* were susceptible to DpnI digestion **(Figure 4B).**

**Figure 4.**
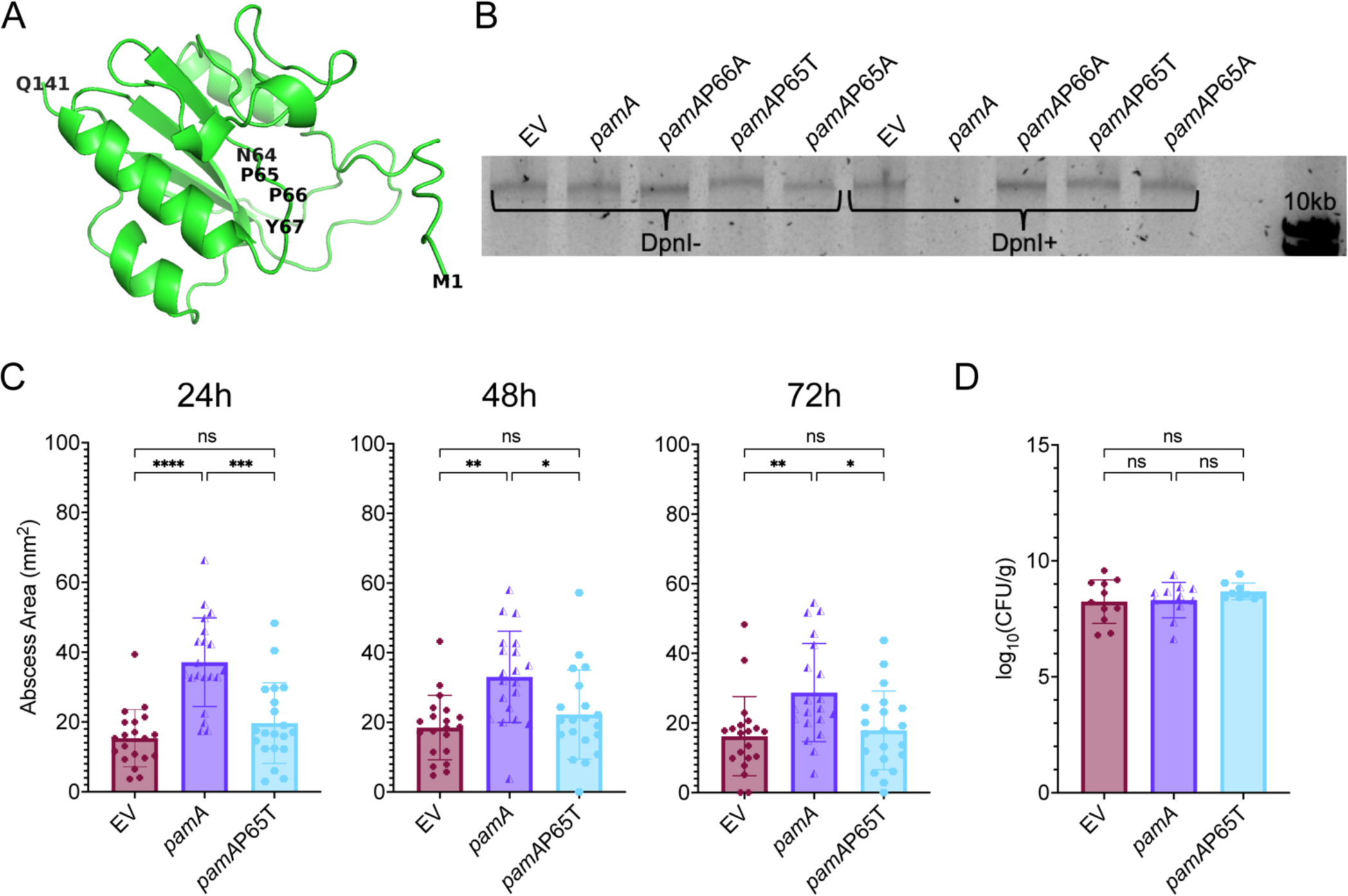
The *pamA*-mediated skin abscess phenotype depends on the methylase activity of PamA. **(A)** Predicted structure of mΦ11 PamA. Amino acid backbone represented in green, with N-terminus (M1), C-terminus (Q141) and putative active site (N64, P65, P66, Y67) highlighted. Generated by AlphaFold, visualized using PyMol Molecular Graphics System, Version 2.5.2 (Schrödinger, LLC). **(B)** Effect of PamA point mutants on methylase activity. Genomic DNA was isolated from LAC* strains containing the indicated *pamA* alleles and digested with DpnI (DpnI+) or PBS control (DpnI-), then visualized on a 1% agarose gel. The analysis confirms that PamA methylates at the predicted G<u>A</u>TC site and that PamA point mutants lack methylation activity. EV, empty vector. **(C)** Skin abscess size. Abscess area of LAC* with EV (maroon, N=20 abscesses, strain RU129), *pamA* (purple, N=20 abscesses, strain RU121), and *pamA*P65T (cyan, N=20 abscesses, strain RU162) at the indicated time points after skin infection with ∼1x10^7^ CFU of bacteria per abscess. Data are pooled from two independent experiments and represent mean ± SD. Statistical significance was determined with the Kruskall-Wallis test and Dunn’s multiple comparisons test, \**P*≤.05, \*\**P*≤.01, \*\*\**P*≤.001, \*\*\*\**P*≤.0001. **(D)** Bacterial burden in abscesses. Skin abscesses from infections in panel C (N=9-11 abscesses per strain) were harvested at 72h and CFU enumerated. Data represent mean ± SD. Statistical significance was determined with the Kruskall-Wallis test and Dunn’s multiple comparisons test.

For in vivo studies, we used the *S. aureus* strain containing *pamA*P65T, since this substitution exhibited the most significant decrease in Dam methylation activity (28). Consistent with the hypothesis that the methylation activity of PamA contributes to the increased abscess size, LAC*::*pamA*P65T produced abscesses that were 32-47% smaller than LAC*::*pamA* abscesses and similar in size to LAC*::EV control **(Figure 4C)**. The P65T amino acid change eliminated the *pamA*-mediated large-size skin abscess size phenotype without affecting tissue bacterial burden in the underlying tissues **(Figure 4D)**. We conclude that the DNA methylation activity of PamA increases abscess virulence.

### Identification of genes involved in *pamA*-mediated virulence

We proceeded to investigate whether *pamA* epigenetically regulates bacterial gene(s) that result in the hyper-abscess phenotype. Notably, *pamA* is constitutively expressed in LAC*::*pamA*, allowing us to bypass the *in vivo* induction needed to produce mΦ11-related phenotypes. Thus, we performed RNA-seq with LAC*::*pamA* and LAC*:EV strains during exponential growth in nutrient restrictive (RPMI) medium chosen to resemble nutrient availability under infectious conditions in human plasma (30). LAC*::*pamA* induced widespread transcriptional changes in CA-MRSA compared to the LAC*::EV control, with 483 genes differentially expressed (232 overexpressed, 250 under-expressed, adjusted *P* <0.05) **(Figure 5A).** The most significantly upregulated gene in LAC*::*pamA* compared to LAC*::EV encodes fibronectin-binding protein A (*fnbA*; FnBPA) **(Figure 5A)**. qRT-PCR confirmed a 15-fold increase in *fnbA* transcription in the LAC*::*pamA* strain compared to LAC*::EV **(Figure 5B)**. FnBPA is a *S. aureus* cell-wall anchored protein that binds adhesive matrix molecules, increasing *S. aureus* invasion into non-professional phagocytic cells (31-33). FnBPA also induces platelet aggregation (34), promotes biofilm formation (35-37), and has been implicated as a virulence factor in endocarditis (38), sepsis (39), implant infections (40), and skin and soft tissue infections (41). Collectively, these observations suggest that *fnbA* plays a role in *pamA*-mediated virulence.

**Figure 5.**
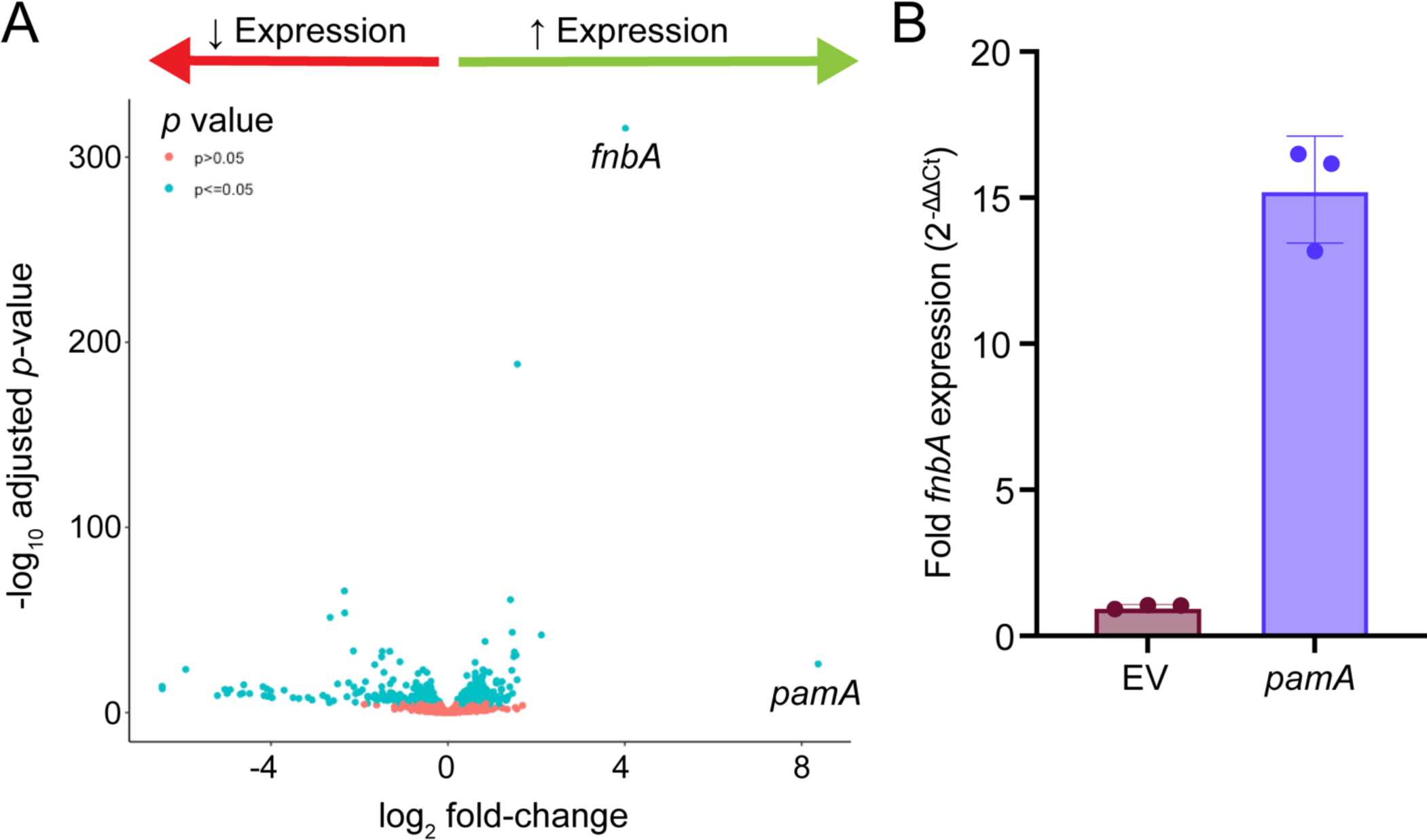
*pamA* induces widespread transcriptional changes including a large increase in the expression of fibronectin-binding protein A (*fnbA;* FnBPA). **(A)** Whole genome transcriptome. Volcano plot of RNA-sequencing data comparing LAC* strains containing *pamA* (N=3 biological replicates, strain RU121) or empty vector (EV) control (N=2 biological replicates, strain RU129) after 5 hours of growth in RPMI media. Data points to the right of zero (green arrow) represent upregulated genes in LAC*::*pamA* and data points to the left of zero (red arrow) represent downregulated genes in LAC*::*pamA*; *pamA* and *fnbA* are highlighted. Blue data points represent genes that achieved statistical significance (*P* ≤0.05); pink data points indicate genes that did not. **(B)** Effect of *pamA* on *fnbA* expression. Quantitative real-time PCR of *fnbA* in LAC* strains containing *pamA* or EV control. Strains were grown and prepared in the same manner as panel A. Data represent mean ± SD of three biological replicates.

### pamA increases biofilm production in vitro and in vivo by increasing FnBPA

The upregulation of *fnbA* expression observed in *pamA* containing strains, coupled with its association with biofilm-related infections (42) suggest that *pamA* increases the formation of biofilms. Indeed, we found that LAC*::*pamA* nearly doubled biofilm production compared to LAC*::EV in an in vitro biofilm assay **(Figure 6A)**. This phenotype reverted to LAC*::EV when *pamA* contained an inactivating point mutation **(Figure 6A).** Thus, the methylase activity of *pamA* increases biofilm production.

**Figure 6.**
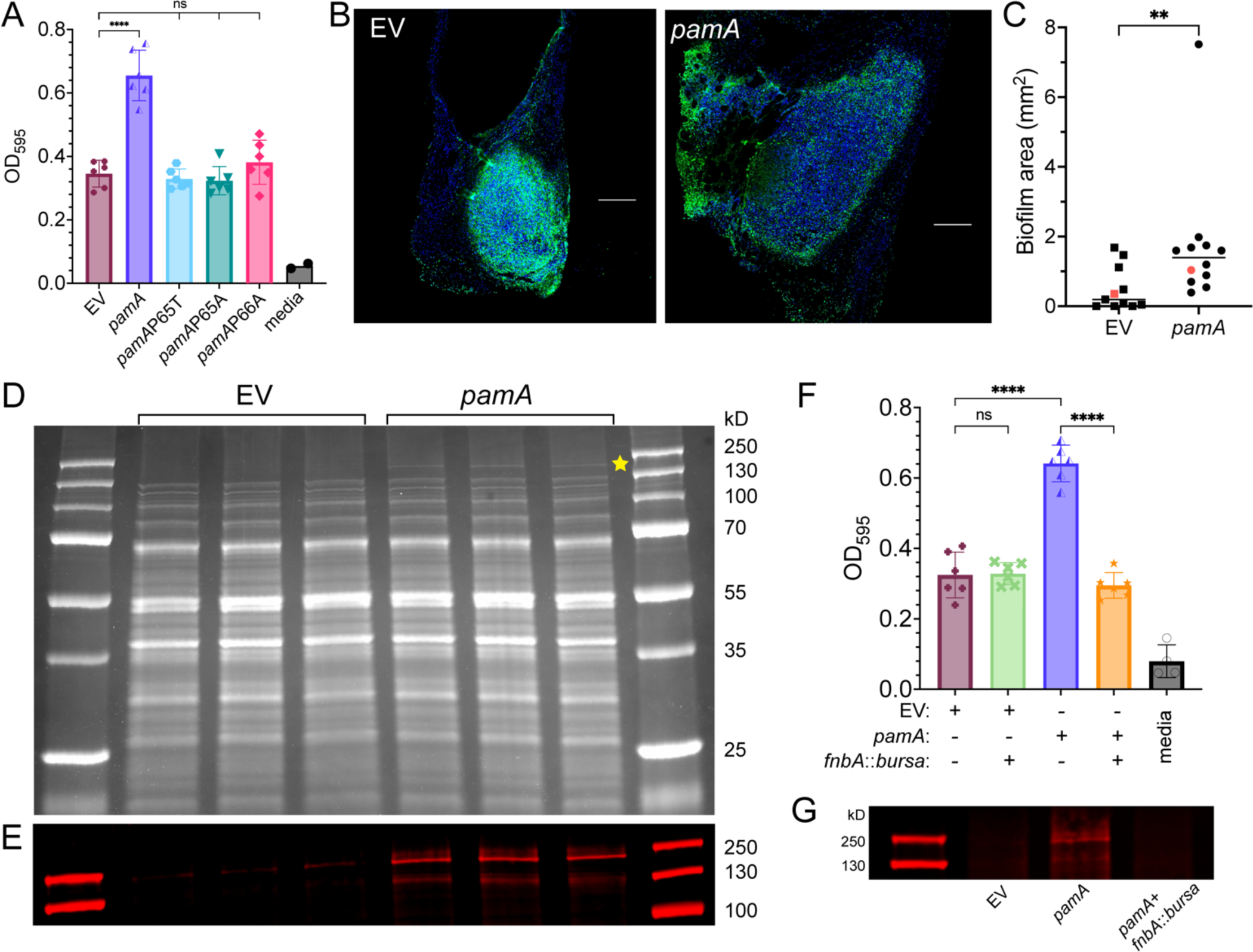
The methylase activity of *pamA* acts to increase biofilm production through fibronectin-binding protein A (*fnbA;* FnBPA). **(A)** Effect of *pamA* methylase activity on biofilm production. In vitro biofilm production by LAC* strains with the indicated *pamA* alleles integrated into the chromosome, quantified by optical density after static growth for 24 h (OD). EV, empty vector control. Data represent mean ± SD of six biological replicates per strain, pooled from two independent experiments. **(B)** Effect of *pamA* on biofilm formation in abscesses. Representative images of skin abscess tissue stained for DAPI (blue) and 5-methylcytosine (5mc, green) 72 h after infection with ∼1x10^7^ CFU of LAC* containing *pamA* (strain RU121) or EV (strain RU129). Scale bar (white) is 200 µm. **(C)** Biofilm area of LAC* containing *pamA* (N=12 abscesses, strain RU121) or EV (N=11 abscesses, strain RU129) quantified as the difference between DAPI and 5mC staining (48). Red data points correspond to representative images in panel B. Data was pooled from two independent experiments. Statistical significance was determined with the Mann-Whitney test, \*\**P*≤.01. **(D)** Cell wall proteins. Cell wall-associated proteins from biofilms of LAC* strains containing *pamA* or EV (three biological replicates each) were separated by SDS-PAGE and stained with Coomassie blue. Gel image is representative of two independent experiments. Yellow star corresponds to the band of interest. **(E)** Identification of FnBPA bands. Western blot of cell wall associated protein bands from panel D using polyclonal anti-FnBPA, focusing on high molecular weight protein band area. (**F)** Biofilm production. In vitro biofilms from LAC* strains containing the indicated genetic changes was quantified by optical density (OD). Data represent mean ± SD of six biological replicates per strain, pooled from two independent experiments. **(G)** FnBPA production. Western blot of cell-wall associated proteins during in vitro biofilm production by the indicated strains.

Biofilm has traditionally been associated with device-related infections (43, 44), endocarditis (45), and osteomyelitis (46). However, we and others have found that biofilms also form during *S. aureus* deep tissue abscess infections (47, 48). Additionally, skin abscess size correlates with *in vitro* biofilm formation with *S. aureus* (49). To determine whether *pamA*-mediated biofilms form *in vivo*, we quantified biofilm production in skin abscess tissue of LAC*::*pamA* and LAC*::EV control strains by immunofluorescent staining of extracellular bacterial DNA (48), an abundant component of *S. aureus* biofilms (50). LAC*::*pamA* strains produced six-fold more biofilm compared to the LAC*::EV control **(Figures 6B-C)**. Thus, *pamA* stimulates biofilm production in skin abscesses, supporting the idea that *pamA*-mediated biofilm production is important for pathogenesis of the skin abscess phenotype.

To test whether FnBPA production was increased in *pamA*-associated biofilms, we compared levels of FnBPA in biofilms from LAC*::*pamA* and LAC*::EV control strains. Bacterial cell-wall-associated proteins from in vitro biofilms, as determined by SDS-PAGE, are shown in **Figure 6D**. A distinct, high molecular weight protein band was observed to be more abundant in LAC*::*pamA* compared to control strain LAC*::EV; there was otherwise considerable similarity in the distributions of the corresponding bands obtained from the two strains. Consistent with our transcriptional data, the high molecular weight protein band was identified as FnBPA by mass spectrometry **(Figure S4)** and confirmed by western blot **(Figure 6E).**

To determine whether FnBPA was responsible for increased biofilm production, we compared biofilm formation in a *fnbA-*inactivated mutant of LAC*::*pamA* (LAC*::*pamA* plus *fnbA*::*bursa*) and a control strain carrying an empty vector (LAC*:EV plus *fnbA*::*bursa*). The results show that LAC*::*pamA* plus *fnbA*::*bursa* phenocopied the biofilm production of the LAC*:EV strain **(Figure 6F);** thus, the *fnbA* inactivation reversed the biofilm-enhancing effect of *pamA*. Western blot of cell wall-associated proteins from biofilm-associated bacteria confirmed that LAC*::*pamA* increased FnBPA production **(Figure 6G).** Thus, *pamA* increases biofilm production in USA300 LAC* by increasing production of FnBPA.

### pamA increases skin infection virulence through fibronectin-binding protein A (fnbA)

To investigate if *fnbA* was responsible for *pamA*-associated skin abscess virulence, we compared LAC*::*pamA* plus *fnbA*::*bursa* mutant and control LAC*::*pamA* strains. Strain LAC*::*pamA* plus *fnbA::bursa* produced 56-62% smaller abscesses than strain LAC*::*pamA* **(Figure 7A).** At the same time, inactivation of *fnbA* did not affect abscess size in the LAC*::EV control, indicating that the upregulation of *fnbA* by *pamA* is required for the observed phenotype. We found no difference in abscess tissue bacterial CFU related to the presence of *pamA* or *fnbA* **(Figure 7B)**, supporting the idea that *fnbA* is necessary for the increased inflammatory response seen in LAC*::*pamA*.

**Figure 7.**
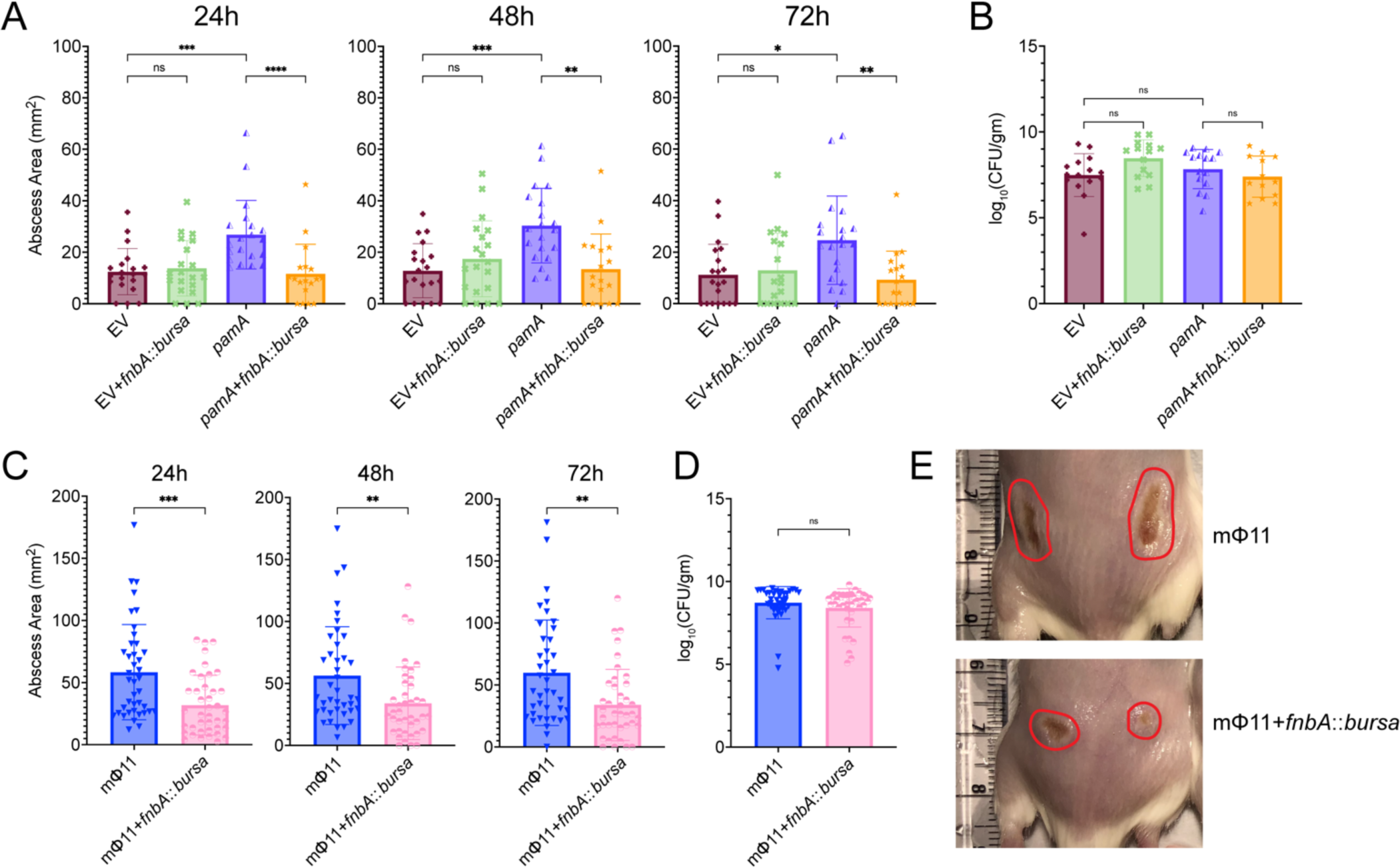
*fnbA* deficiency reverses the *pamA*-mediated skin abscess phenotype. **(A)** Effect of *fnbA* on the *pamA*-mediated skin abscess phenotype. Abscess area of LAC* with EV control (maroon, N=20 abscesses, strain RU129), EV+*fnbA::bursa* (green, N=18-20 abscesses, strain RU170), *pamA* (purple, N=20 abscesses, strain RU121), or *pamA*+*fnbA::bursa* (orange, N=18 abscesses, strain RU169) after skin infection with ∼1x10^7^ CFU of bacteria per abscess for the indicated times. Data are pooled from two independent experiments and represent mean ± SD. Statistical significance was determined with the Kruskall-Wallis test and Dunn’s multiple comparisons test, \**P*≤.05, \*\**P*≤.01, \*\*\**P*≤.001, \*\*\*\**P*≤.0001. **(B)** Bacterial burden in abscesses. Skin abscesses in panel A (N=14-15 abscesses per strain) were harvested for CFU enumeration 72 h post-infection. Data represent mean ± SD. Statistical significance was determined with the Kruskall-Wallis test and Dunn’s multiple comparisons test. **(C)** Effect of *fnbA* on the mΦ11-mediated skin abscess phenotype; transposon insertion in *fnbA* in mΦ11-containing strains confirms that *fnbA* is necessary for the skin abscess phenotype. Abscess area of LAC* containing prophage mΦ11 (blue, N=40 abscesses, strain BS989) and mΦ11+*fnbA::bursa* (pink, N=40 abscesses, strain RU171) after infection with ∼1x10^7^ CFU of bacteria per abscess for the indicated times. Data are pooled from four independent experiments and represent mean ± SD. Statistical significance was determined with the Mann-Whitney test, \**P*≤.05, \*\**P*≤.01, \*\*\**P*≤.001, \*\*\*\**P*≤.0001. **(D)** Skin abscesses from infections in panel C were harvested at 72 h post-infection and CFU enumerated. Data represent mean ± SD. Statistical significance was determined with the Kruskall-Wallis test and Dunn’s multiple comparisons test. **(E)** Representative images at 72 hours post-infection of the indicated strains from panel E, abscess area circled in red.

To ensure that the in vivo phenotype is specific to *pamA*, and not an artifact of *pamA* overexpression, we compared mice infected with an *fnbA-*inactivated mutant of mΦ11 (LAC*/mΦ11:*fnbA*::*bursa*) to those infected with control strain LAC*/mΦ11. The results show that *fnbA* is critical for the increased virulence observed with mΦ11 containing strains **(Figures 7C-E)**, indicating that overexpression alone cannot explain the increased skin infection virulence demonstrated in LAC*::*pamA*. We conclude that LAC*::*pamA* can provide insights into the role of *pamA* in the virulence of mΦ11-containing strains and that *fnbA* is an essential downstream virulence factor in the *pamA* regulatory pathway underlying increased virulence during skin infection.

## Discussion

The present findings, in conjunction with previous molecular epidemiology observations (5), indicate that the mechanism by which mΦ11 primed the epidemic USA300 Brooklyn clone for success is via a phage adenine methyltransferase (*pamA*) that increases bacterial virulence during skin abscess infection. We discovered that *pamA* mediates this increase by rewiring the strain through epigenetic modifications that induce expression of *fnbA*, a virulence factor essential for the enhanced virulence of mΦ11-containing USA300. Interestingly, we found that *fnbA* is not a skin abscess virulence factor in the absence of *pamA*. FnBPA is, in turn, associated with increased abscess inflammation and an increased ability to form biofilms. Successful spread of the Brooklyn clone after acquisition of mΦ11 is tied to subsequent selection of antimicrobial resistance (5). Thus, analysis of the USA300-phage interaction in the context of infection identified an unappreciated role for epigenetics in the evolution of virulence in *S. aureus* and ultimately antimicrobial resistance in patients.

Recent reports link the *E. coli* prophage Φstx104-encoded methyltransferase (M.EcoGIII) to global transcriptional changes in a hypervirulent strain O101:H4 variant that is associated with an outbreak of hemolytic uremic syndrome (HUS) (26). Thus, the phenomenon of prophage methyltransferase acquisition in epidemic strains may be common. However, the contribution of M.EcoGIII transcriptional regulation to virulence of *E. coli* O101:H4 is confounded by the observation that ΦStx104 encodes Shiga toxin genes *stxA* and *B* that cause HUS. To our knowledge, mΦ11 is the only example in which differential regulation of a core virulence factor by a phage-encoded methyltransferase is sufficient for elevated virulence during infection.

To date, studies on the role of prophage in pathogenesis focus largely on the role of prophage-encoded secreted toxins and immune modulators (51). Nevertheless, many, if not most, prophages lack known virulence or fitness factors (12) and frequently contain orphan methyltransferases (52). Thus, our findings, and those in *E. coli*, support the idea that prophage-mediated regulation of host bacterial gene expression may be a critical feature of the bacterium-phage interaction. In this connection, we note that ΦN315 phage-encoded *sprD*, a regulatory RNA, can enhance virulence in *S. aureus* (53, 54). This finding suggests the existence of alternative routes to hypervirulence with phage that involves gene regulation.

Our results, and those of M.EcoGIII in *E. coli,* also support the idea that DNA methylation has implications beyond that of bacterial defense against foreign DNA. Recent work showed that horizontal acquisition of phage-encoded methyltransferases and associated changes in gene expression are linked to speciation in *V. cholera* (55) and *M. abscessus* (56). In contrast, although restriction-modification systems are well characterized in *S. aureus* (reviewed in (57)), little has been known about the role of methyltransferases in staphylococcal gene regulation. Thus, the present study forms a knowledge base whereupon the role of methyltransferase in the pathogenesis of staphylococcal disease can now be interrogated by genome-wide mapping to assess the role of DNA modification events in staphylococcal virulence.

Two additional comments are relevant to the work described above. First, our present and prior (5) findings that mΦ11 phenotypes are specific to in vivo analyses supports the notion that in vitro analyses of virulence genes and regulators do not necessarily correlate with virulence during infection (58). This may be especially true for phage-mediated virulence, where induction is often critical to stimulate expression of phage-encoded genes (59). Therefore, future investigations to determine prophage effects on virulence should prioritize in vivo models. Second, the observation that biofilm is associated with virulence during skin abscesses caused by USA300 is concordant with a prior study linking biofilm and deep tissue abscesses (41). Accordingly, we suggest that skin infection should be included in the established MRSA infectious syndromes (e.g. device infections, endocarditis, osteomyelitis) with which biofilm is traditionally associated.

In summary, our report identifies molecular mechanisms underlying the relationship between prophage, virulence, and the emergence of an adapted CA-MRSA clone. Several questions concerning the role of phage-encoded methyltransferases are raised by the present study. For example, what is the host tissue-specific signal in vivo and does it require phage (or phage gene) induction? Moreover, how does *pamA* alter methylation patterns and what specific difference in methylation is responsible for increased *fnbA* expression and the enlarged abscess phenotype? Third, is the phage-encoded methyltransferase mechanism of gene regulation useful in other ecological contexts, such as colonization? And, lastly, what is the frequency and distribution of orphan methyltransferase in natural populations of phage and *S. aureus*? Implicit in these questions is the view that phage are independently evolving entities that have acquired pathogenesis-adaptive genes. This behavior optimizes their existence as parasites, which they have fine-tuned through multiple functions that enhance the fitness of their bacterial hosts.

## Methods

### Bacterial strains and growth conditions

Bacterial strains, plasmids, and oligonucleotides used in this study are described in **Table 1**. *S. aureus* colony formation was on 5% sheep blood agar (SBA) or tryptic soy agar (TSA) plates and *E. coli* was on Luria Bertani (LB) plates. *S. aureus* strains were grown in tryptic soy broth (TSB) medium at 37°C with orbital shaking at 180 rpm. Plates and media were supplemented with selective antibiotics when appropriate [ampicillin (Amp) 100 µg/mL, chloramphenicol (Cm) 10 µg/mL, erythromycin 5 µg/mL (Erm) or CdCl_2_ (Cd) 0.4 mM unless stated otherwise]. Transductions were performed with phage 80α using established protocols (60); transductants were selected on TSA plates with appropriate antibiotics. PCR amplifications used Phusion^TM^ Plus PCR Master Mix (Thermo Scientific, #F631) and oligonucleotides from Integrated DNA Technologies (IDT). Detailed strain construction methods are provided in **Supplemental Method A**. Briefly:

**Table 1:**
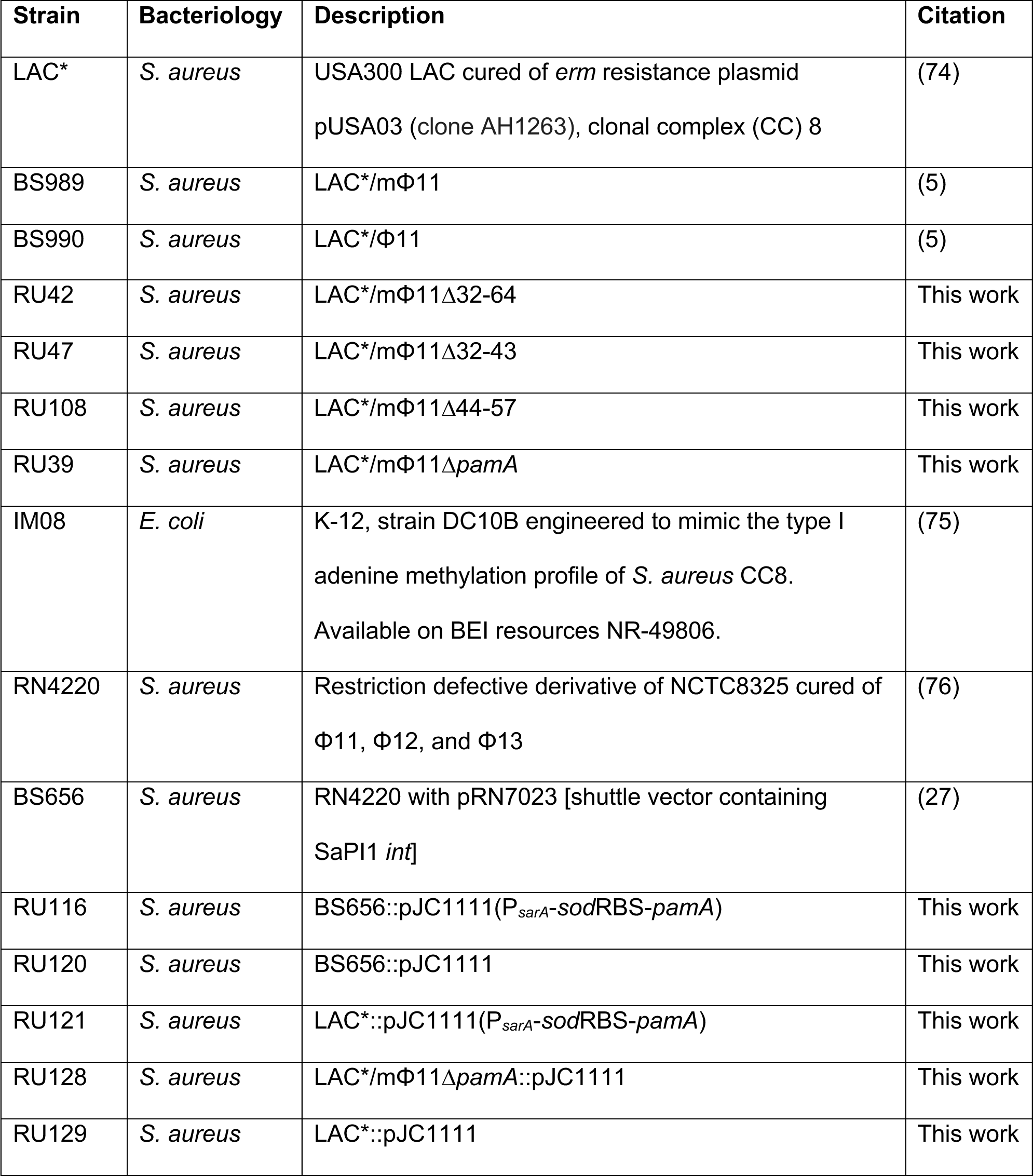
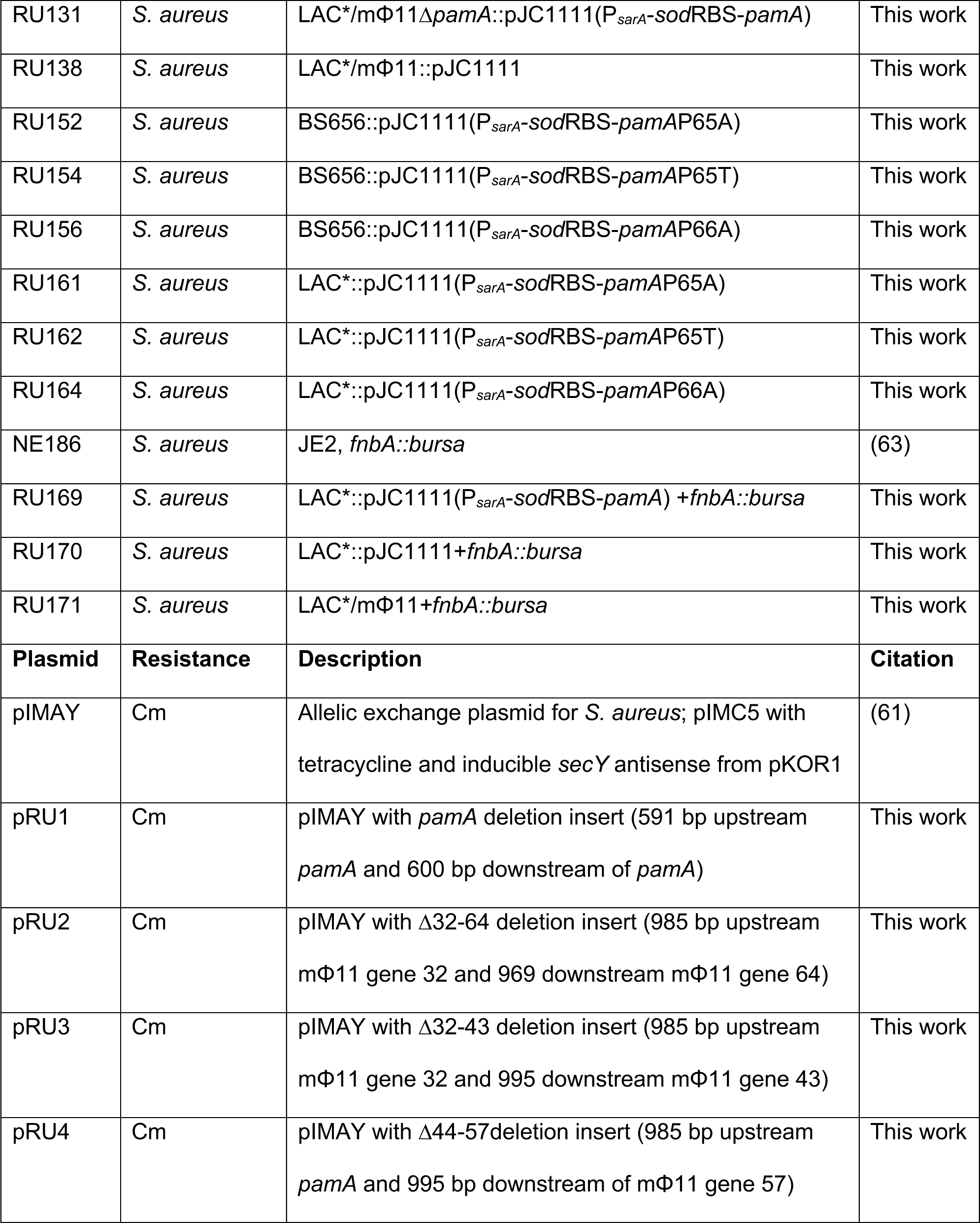
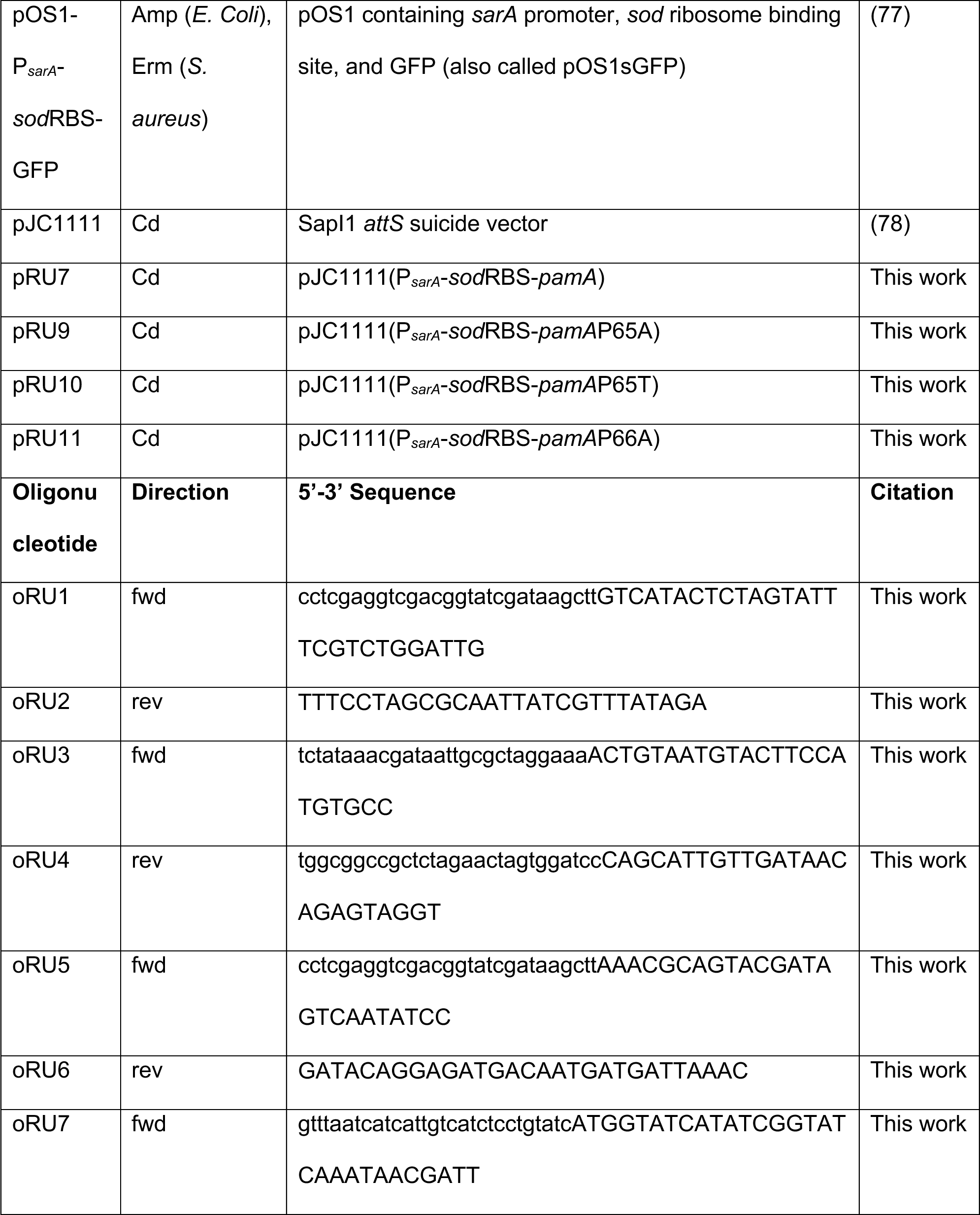
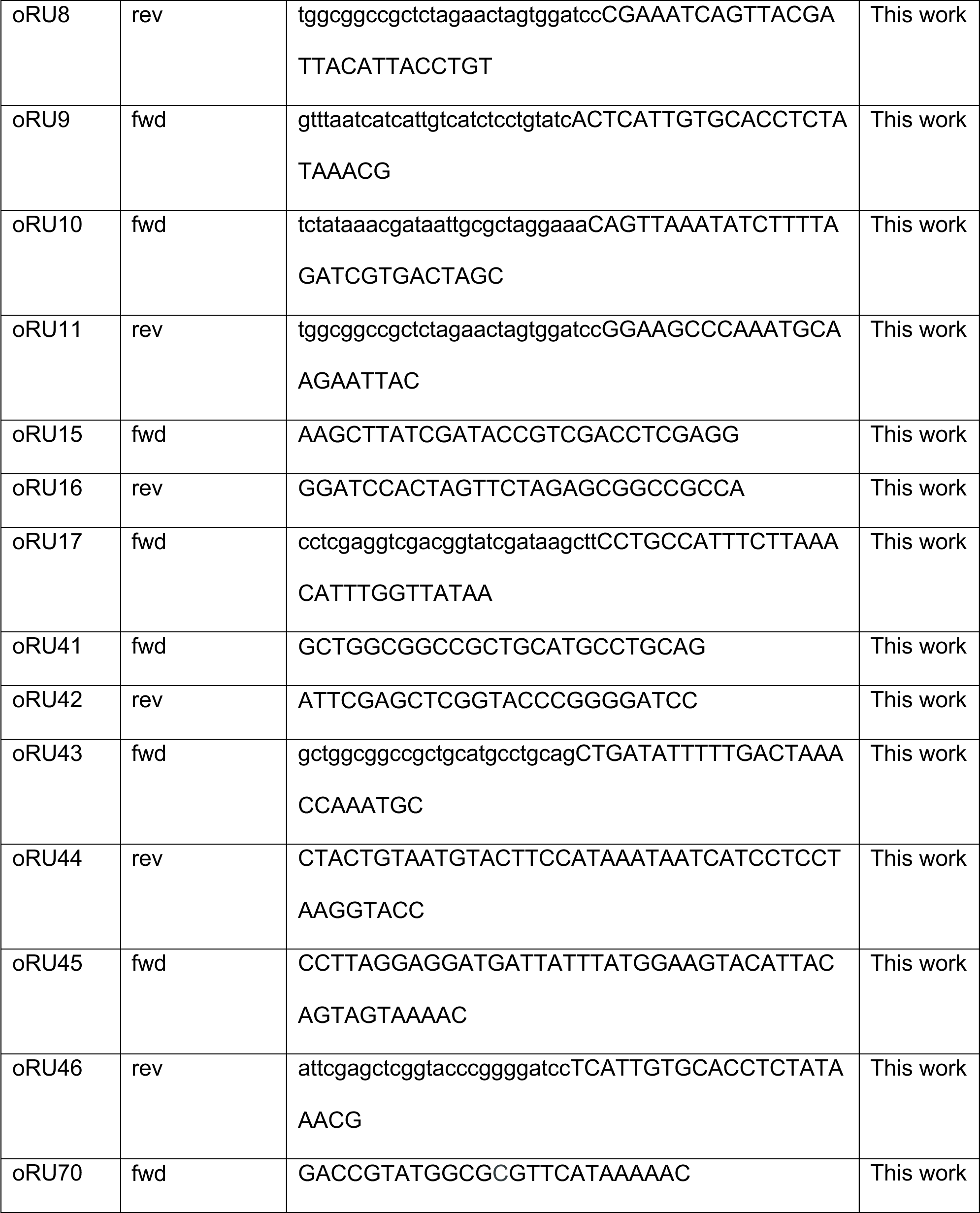
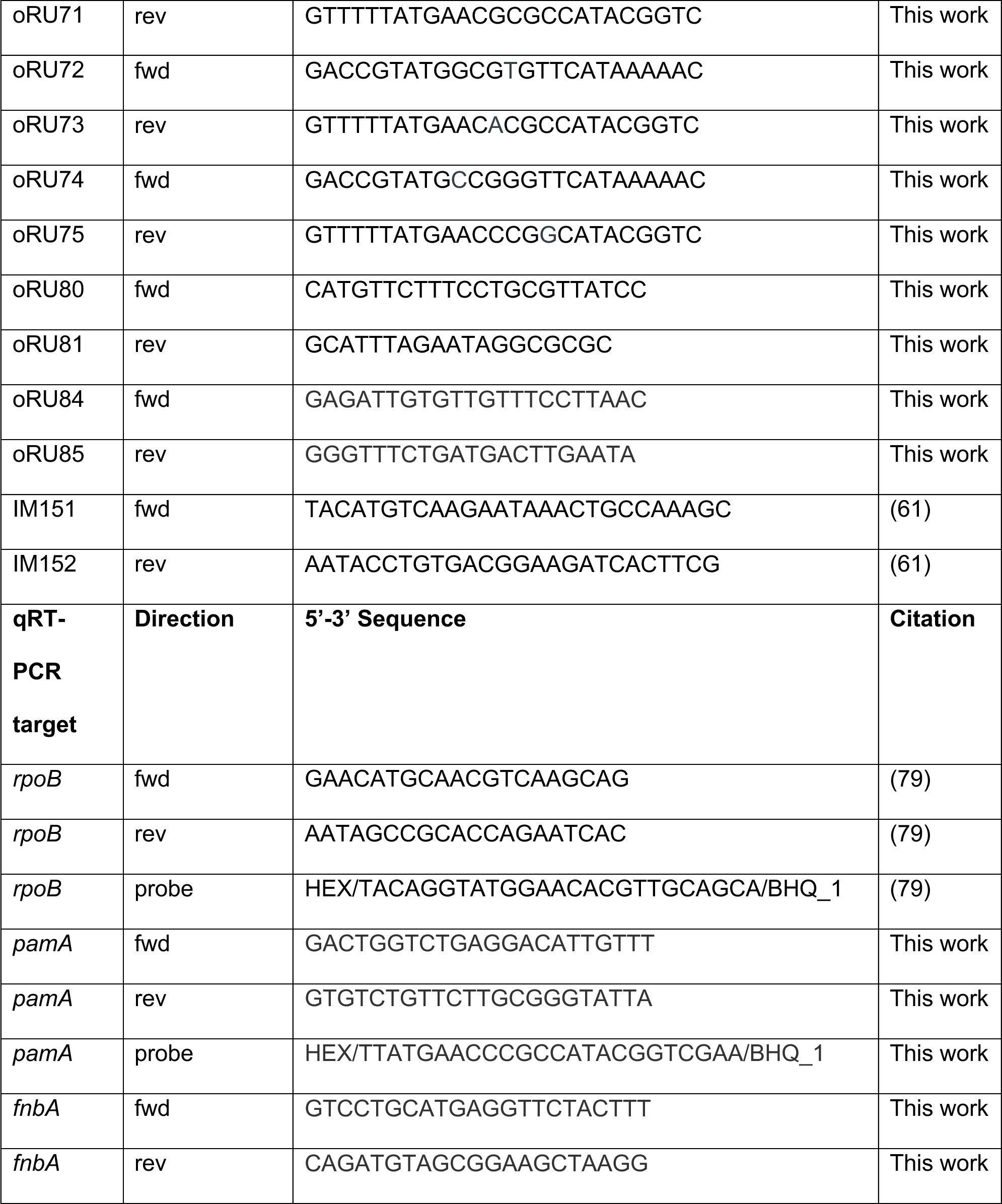
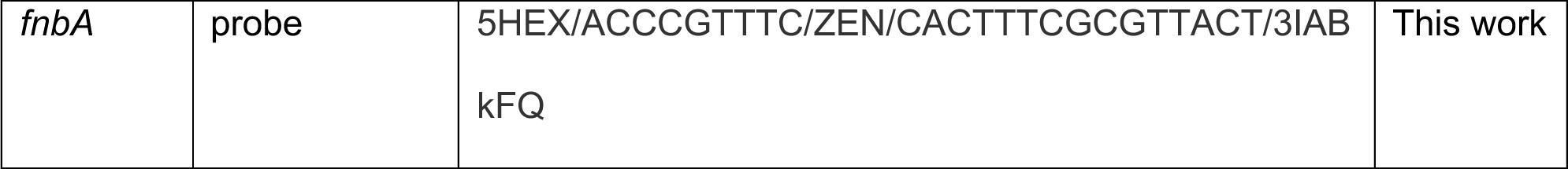
Strains, Plasmids, Oligonucleotides, and Probes Used in this Study.

Strains containing in-frame deletions (RU39, RU42, RU47, and RU108) were engineered in strain LAC*/mΦ11 (BS989) by allelic exchange with cloning plasmid pIMAY (61). Deletions were confirmed by Sanger sequencing (Psomagen,Inc.) and comparative sequence analysis **(Figures S2, S3)**, as outlined below.

Strains containing a single-copy chromosomal insertion of a constitutively expressed *pamA* (P*_sarA_*-*sod*RBS-*pamA*) or empty vector (EV) were generated by insertion of pRU7 or pJC1111, respectively, at the *S. aureus* pathogenicity island 1 (SapI1) site of strain BS656 (27), then transducing the mutation with phage 80α into LAC* (resulting in strains RU121 and RU131) or LAC*/mΦ11Δ*pamA* (resulting in strains RU129 and RU128). The presence and location of the P*_sarA_*-*sod*RBS-*pamA* and EV inserts were confirmed by PCR.

To construct strains with inactivating *pamA* point mutations, complementary oligonucleotides that contained the desired *pamA* mutation were used to amplify P*_sarA_*-*sod*RBS-*pamA* from pRU7 template DNA, then the amplification products were sewn together by overlap extension PCR (62). P*_sarA_*-*sod*RBS-*pamA*P65A, P*_sarA_*-*sod*RBS-*pamA*P65T and P*_sarA_*-*sod*RBS-*pamA*P66A fragments were inserted into pJC1111 with Gibson assembly resulting in pRU8, pRU9 and pRU10, respectively. pRU8, pRU9, and pRU10 were transformed into competent *E. Coli* DH5⍺ (New England Biolabs, #C2987H) per manufacturer instructions, electroporated into BS656 for insertion into the SapI1 *att* site (27), and transduced into LAC* with phage 80α, resulting in strains RU161, RU162, and RU164, respectively. Point mutations were confirmed by Sanger sequencing.

To construct strains with *fnbA::bursa* transposon insertions, phage 80α lysate of strain NE186 (*fnbA*::*bursa,* Erm^r^) (63) was used to transduce RU121, RU129, and BS989, generating RU169, RU170, and RU171, respectively. PCR amplification across the *fnbA::bursa* insertion site was performed to confirm the transposon insertion.

### Mapping deletions using whole genome sequencing (WGS)

Extracted purified gDNA was quantified with the Quant-it picogreen dsDNA assay kit (Invitrogen, #P7589) prior to library prep. Samples were normalized by concentration and libraries prepared with the Illumina DNA prep (M) Tagmentation kit (#20018705). Each library was combined equimolar and sequenced as paired-end 150-bp reads using the Illumina Novaseq 6000 system with the S1 300 cartridge and flow cell.

For WGS analysis, BWA v0.7.17 (64) was used to map the raw short-read sequences of the *S. aureus* samples (strains LAC*, BS989, RU39, RU42, RU47, and RU108) to a *S. aureus* reference assembly (NCBI accession GCF_015475575) and the mΦ11 phage (GenBank accession PP554657), resulting in one alignment file per sample. The depth of mapped reads was computed with bedtools v2.30.0 (65) using the command bedtools genomecov -iBAM file.bam -d, where file.bam stands for each of the sample alignment files. This output files tabulating contig name, start site, end site, and number of reads covering each base. Read coverage per base were loaded into R v4.2.0 (R Core Team, 2021) for visualization. Plots were created with ggplot2 (66).

### Growth Curves

Overnight cultures were diluted (1:1000) into fresh TSB or RPMI medium (Sigma-Aldrich, #R6504) and growth was monitored at 37°C in 100-well (150 µL-well) honeycomb plates (Thermo Scientific, #12871511), using a Bioscreen C Analyzer (Thermo Labsystems), measuring OD_600_ at 30-min intervals. The curves represent averaged values from three biological samples. Each biological sample was run as ten technical replicates.

### Secreted protein preparation

Overnight cultures were pelleted, washed with sterile PBS, OD normalized, and diluted 1:200 into TSB medium for growth at 37°C shaking at 180 rpm. After 6 h and 24 h, cells were centrifuged at ∼3,200 x g for 15 minutes to remove bacteria and aliquots (1.3 mL) of supernatant were passed through 0.2 µM filter (Thermo Scientific, #723-2520). The supernatants were precipitated with 100% trichloroacetic acid using established protocols (67).

### Animal Infections

Five-week-old female Swiss Webster mice (Envigo) were anesthetized with Avertin (2,2,2-tribromoethanol dissolved in tert-Amyl alcohol and diluted in sterile PBS to a final concentration of 2.5% vol/vol) intraperitoneal injection (300 µL). Mice were shaved with mechanical clippers and ∼1x10^7^ CFU of bacteria was injected (100 µL) subcutaneously into each flank using Adson forceps (68). For daily abscess measurements, mice were briefly anesthetized with inhaled isoflurane. Abscess diameter was measured with digital calipers (Thermo Scientific, #14-648-17). Abscess area was quantified using digital photography and ImageJ (69). Measurements were standardized to a centimeter ruler in-frame. At 72 h post-infection, mice were euthanized, and abscesses excised with an 8mm punch biopsy (Integra Life Sciences). Tissue biopsies were either prepared for histology as described below or homogenized for CFU enumeration and/or cytokine analysis. If both CFU enumeration and histology or other preparation was required, left flank biopsies were homogenized for CFU enumeration while right flank biopsies were used for the additional analysis, to minimize bias. For homogenization, biopsy samples were added to 2 mL conical screw cap tubes (Thermo Scientific, #023-681-344 and #02-681-358) with sterile PBS (1mL) and a single 0.25” ceramic sphere (MP Biomedicals, #116540034), weighed, and homogenized by three cycles in a FastPrep-24 homogenizer (MP Biomedicals) at four m/s for 60 s. Homogenates were serially diluted in sterile PBS and plated on TSA for CFU enumeration. For cytokine analysis, 1x Halt protease inhibitor cocktail (Thermo Scientific, #78429) was added to homogenates and samples were stored at -80°C.

### Histology

Skin biopsies were immobilized in cassettes (Simport Scientific, #M490-2), fixed in 10% formalin for 72 h at 4°C, washed in sterile PBS three times for 20 min, then dehydrated with increasing concentrations of ethyl alcohol (EtOH) before storage in 70% EtOH at 4°C. Fixed and dehydrated specimens were embedded in paraffin and 5 µm sections were performed through the center of the abscess for H&E and gram staining. Slides were scored for inflammatory burden (mild/moderate/severe) and abscess architecture (nodular/diffuse) by a board-certified dermatopathologist (RK), who was blinded to the sample identity throughout.

### Cytokine analysis

Skin abscess cytokine profiles were obtained using the MILLIPLEX MAP Mouse Cytokine/Chemokine Magnetic Bead Panel (Millipore Sigma, MCYTMAG-70K-PX32). Samples were prepared as per manufacturer’s instructions. Data were acquired using a Luminex MABPIX instrument and analyzed using xPONENT software (Millipore). Statistical analyses were performed for each individual cytokine.

### DpnI digestion

gDNA was extracted from strains RU121, RU129, RU161, RU162, and RU164, digested using DpnI (New England Biolabs, #R0176S) per manufacturer protocol, separated on a 1% agarose gel containing SYBERSafe (Thermo Scientific, #S33102) and imaged in a Chemidoc Imager (Bio-Rad Laboratories).

### RNA preparation and sequencing

For transcriptional profiling of strains LAC* and LAC*/m⏀11 **(Figure S1E)**, two independent overnight cultures were diluted (1:100) into fresh TSB medium (5 mL) and grown at 37°C shaking at 180 rpm to early (3 h) or late exponential growth phase (6 h). For transcriptional profiling of strains LAC*::EV and LAC*::*pamA* **(Figures 5A-B),** three independent overnight cultures were diluted (1:100) into RPMI medium (15 mL) and incubated at 37°C with shaking at 180 rpm to exponential growth phase (5 h).

For RNA extraction, cells were concentrated by centrifugation (3,400 x *g* for five min), resuspended in 1 mL Trizol (Invitrogen, #15596026), and disrupted using lysis matrix B (MP Biomedicals, #116911050) tubes in a FastPrep-24 (MP Bio) at six m/s, for 30s, three times. Samples were centrifuged at 12,000 x *g* for 10 min at 4°C and the upper phase was transferred into a new RNA-free tube containing ice cold Trizol (500 µL), gently mixed, and incubated (five min) at room temperature, then chloroform (200 µL) was added and samples were centrifuged at 12,000 x *g* for 15 min at 4°C. The aqueous phase was mixed with isopropanol (500 µL) and transferred to RNeasy column (Qiagen #74004) for washing and RNA elution. RNA was visualized on the Agilent 2100 Bioanalyzer system using a Bioanalyzer Nanochip run with the Prokaryote setting. Libraries were prepared with total RNA (500 ng per sample) of the high-quality samples (RNA integrity number 9-10) using the Illumina stranded Total RNA Prep, Ligation with Ribo-Zero Plus kit (#20040529) per manufacturer’s instructions. PCR Amplification was run with 11 total cycles. The libraries were visualized on the Agilent 4200 Tapestation System and concentration was quantified by Qubit (Thermo Scientific). Libraries were pooled equimolar and sequenced as paired-end 50 bases on the Illumina Novaseq 6000 system on one lane of the SP 100 cycle flow cell kit.

### RNA sequencing analysis

For RNA sequencing of strains LAC* and LAC*/m⏀11 **(Figure S1E),** we used previously established analysis methods (70), with the addition of the mΦ11 sequence to the reference genome. For RNA sequencing of strains LAC*::EV and LAC*::*pamA* **(Figure 5A),** we created a reference assembly by appending the pJC1111 sequence to the AH-LAC assembly (NCBI accession number GCF_015475575.1) and annotating with NCBI Prokaryotic Genome Annotation Pipeline (PGAP) (71). However, the sequence of *pamA* was too short for processing by PGAP, so we manually added it to the gff and genbank annotation files that PGAP produced. We used Bowtie2 v2.4.1 (Langmead & Salzberg, 2012) to align the raw short-read sequences of LAC*::*pamA* and LAC*::EV to the reference assembly. Using the alignment files generated for each sample, the *featureCounts* command in Subread v2.0.1 (Liao et al., 2014) was used to count the reads mapping to each gene in the reference. Read counts per gene and sample were loaded into R v4.2.0 (R Core Team, 2021) for differential expression analysis using the package DESeq2 v1.36.0 (Love *et al.,* 2014). The function DESeq with default settings was used to normalize for library size differences, to estimate dispersion, and to fit negative binomial generalized least squares (GLM) models for each gene. Differential expression testing was performed using the Wald test as implemented by DESeq2. The resulting p values were adjusted using a false discovery rate (FDR) of 10%.

### Quantitative Reverse Transcriptase PCR (qRT-PCR)

RNA was isolated from LAC*::*pamA* and LAC*::EV at exponential growth as described above. DNA was removed with Turbo DNase DNA free kit (Invitrogen, #AM2238), and cDNA was synthesized using the Superscript III First-Strand Synthesis System (Invitrogen, #18080051). qRT-PCR was performed using TaqMan^TM^ Universal PCR Master Mix (Thermo Scientific, #4304437) and primers/probes (IDT) specific to *pamA, fnbA,* and *rpoB*. Three independent biological samples of each strain were run in duplicate and *rpoB* was used to normalize gene expression. Settings on the C1000 CFX96 machine (Bio-Rad Laboratories) were as follows: 50°C for two min, 95°C for 10 min, then 40 cycles [95°C for 15s and 60°C for 1 min]. 2^-△△Ct^ method was used to calculate the relative fold gene expression (72).

### In vitro biofilm assays

Overnight broth cultures of were diluted (1:100) into fresh TSB medium supplemented with 0.25% glucose (TSBG), aliquoted into 96-well (200 µL-well) tissue-culture treated polystyrene plates (Corning, #CLS3799), and incubated statically at 37°C for 24 h. Supernatants were discarded and adherent biofilms were washed three times with sterile PBS (200 µL), fixed with 100% ethanol (200 µL), and stained with crystal violet 0.1% w/v (200 µL) at RT for 15 min. Residual stain was discarded and biofilms were washed three times. Crystal violet was eluted with 33% acetic acid (200 µL) incubated for 10 min, then samples were diluted (1:4) in PBS and quantified by measuring OD_595_ using a Synergy neo2 plate reader (BioTek).

### Biofilm cell-wall associated protein preparation

Overnight cultures were diluted (1:100) into fresh TSBG, aliquoted into six-well (3.6 mL) tissue-culture treated polystyrene plates (Corning, #CLS3516), and incubated statically for at 37°C for 24 h. Supernatants were discarded, biofilms were washed two times with 3.6 mL sterile PBS, resuspended in 1 mL sterile PBS, normalized to OD_600_, and centrifuged (12,000 x *g* for two min). Biofilm pellets were washed twice with PBS (1 mL), re-suspended in a 48 µL mixture of lysostaphin (20 µg/mL), 1x Halt Protease Inhibitor in TSM buffer (10 mM MgCl_2_ 500 mM Sucrose in 50 mM Tris, pH7.5), and incubated for 30 min at 37°C. Samples were centrifuged (12,000 x *g* for two min) and the supernatant (36 µL) was mixed with 4x SDS (12 µL) sample buffer [200mM Tris-Cl (pH 6.8), 588mM beta-mercaptoethanol, 8% SDS, 0.08% bromophenol blue, 40% glycerol, 50mM EDTA]. Samples were boiled for 10 min and stored at -80C.

### Coomassie staining and Immunoblotting

Proteins were separated by sodium dodecyl sulfate polyacrylamide gel electrophoresis SDS-PAGE (12% gel; Bio-Rad Laboratories, #4561043), visualized using InstantBlue Coomassie dye (Abcam, #50-196-3787), and transferred to nitrocellulose membrane for analysis by immunoblot. The membrane was incubated in Everyblot Blocking Buffer (Bio-rad Laboratories, #12010020) for five min blocking at RT, then primary antibody (1:2000 dilution of anti-FnBPA antibody, Abnova, #PAB16068) overnight at 4°C, then secondary antibody (1:25,000 dilution of Alexa Flour 680-conjugated goat anti-rabbit IgG, Invitrogen, #A21076) for one hour at RT. Images were acquired with the Odyssey Clx imaging system and Image Studio software (Li-Cor Biosciences).

### Protein identification by mass spectrometry

High molecular weight bands noted on the LAC*::*pamA* biofilm cell-wall associated protein preparation **(Figure 6D)** were manually excised from gel lanes and stored in 1 mL 1% acetic acid. As a control, similar high molecular weight areas in LAC*::EV lanes and one PBS control lane were excised in the same manner. The proteins were in-gel digested using trypsin as previously described (73). Sample processing, mass spectrometry, and data analysis was performed as described in **Supplemental Method B.**

### Quantification of abscess tissue biofilm by immunohistochemistry

The following methods were adapted from prior work (48). Skin biopsies were fixed with periodate-lysine-paraformaldehyde buffer overnight at 4°C, dehydrated in sucrose (30%) for 24 hours, and frozen in optimum cutting temperature compound (Themo Scientific, #1437365). For *S. aureus* staining, 10 µm-thick skin sections were incubated in bovine serum albumin (2%; BSA) in Tris-buffered saline (TBS) with the primary antibody [1:400 dilution of rabbit anti-*S. aureus* (Abcam, #20920)] at 4°C overnight. The sections were washed three times with 1% BSA in TBS and incubated with the secondary antibody [1:500 dilution of goat anti-rabbit IgG-AF488 (Invitrogen, #A-1108)] and DAPI at 4°C for 1 h. For 5-methylcytosine (5-mC) staining, 10 um-thick skin sections were permeabilized with hydrogen chloride (1.5M, Fisher Chemical) to allow the 5-mC antibody to stain the biofilm. The sections were washed twice with PBS and incubated in 2% BSA plus TBS with primary antibody [1:80 dilution rabbit anti-5-mC antibody (Cell Signaling Technology, #D3S2Z) at 4°C overnight. The sections were washed three times with 1% BSA in TBS and incubated with secondary antibody [1:500 dilution goat anti-rabbit IgG-AF488] and DAPI at 4°C for 1 h. The sections were again washed three times with 1% BSA in TBS and then mounted with cover glass over tissue sections using ProLong Diamond Antifade Mountant (Invitrogen). All of the antibodies were diluted in blocking solution. Imaging and analysis were performed as previously described (48), with the exception that thresholds of 5-mC and DAPI fluorescence intensity were determined based on the staining from mock (sterile PBS) infected skin biopsies.

### Statistics

For comparisons of two groups, normality was determined using the Shapiro-Wilk test. If data were normally distributed, unpaired t-tests were performed. If data were not normally distributed, Mann-Whitney test was used. For comparison of more than two groups, if normality was determined, ANOVA with Tukey’s multiple comparisons test was used. If any group was not non-normally distributed, Kruskal-Wallis test with multiple comparisons was performed. Analyses were performed using Prism version 9.4.1 for Macintosh (GraphPad Software, www.graphpad.com).

### Data Availability

Values for all data points in graphs are reported in the Supporting Data Values file. RNA-sequencing files are deposited in NCBI GEO, accession GSE255351 (corresponding to **Figure S1**) and GSE252862 (corresponding to **Figure 5A**). Whole genome sequencing of *S. aureus* strains constructed during this study are deposited in NCBI SRA database, accession PRJNA1090089. The sequence of prophage mΦ11 can be accessed as GenBank accession PP554657. Additional data will be made available upon request.

### Study Approval

All animal experiments were reviewed and approved by the Institutional Animal Care and Use Committee (IACUC protocol #107203) of New York University Langone Medical Center (NYULMC). All experiments were performed according to NIH guidelines and U.S. federal law.

## Author Contributions

RJU, BS, and VJT designed the study; RJU, MP, RT, II, KL, DB, SMM, SM, TK, EEZ, CZ, and RK performed experiments and generated data; RJU, RT, SM, GP, NS, AP, and HVB analyzed data; RJU, BS, and VJT secured funding; RJU wrote the initial draft. All authors contributed substantial revisions and approved the final version of the manuscript.

## Supporting information

Supporting Data Values file

## Acknowledgements

We thank Francois-Xavier Stubbe at University of Namur for assistance with bioinformatic data analysis; Adam J. Ratner, Andrew J. Darwin, and Jeffery N. Weiser for assistance with experimental design; Boris Reizis for mentorship, resources, and support; John Chen at National University of Singapore for helpful discussions during the project design; The NYU Langone Health Genome Technology Center (RRID:SCR_017929), Experimental Pathology Research Laboratory (RRID:SCR_017928), and Beatrix Ueberheide and the NYU Langone Health Proteomics Laboratory (RRID:SCR_017926) with support from NIH Shared Instrumentation Grant 1S10OD010582-01A1. This work was supported by the National Institute of Allergy and Infectious Diseases at the NIH K08AI163457 (RJU), 1TL1TR001447 (RJU), R01AI137336 and R01AI140754 (BS and VJT), The New York State Department of Health ECRIP (RJU), Pew Latin American Fellows Program (DB), and the National Institute of Arthritis and Musculoskeletal and Skin Diseases at the NIH T32AR064184 (TKK). The content is solely the responsibility of the authors and does not necessarily represent the official views of the NIH. The *fnbA::bursa* mutant strains were created from NE186 of the Nebraska transposon mutant library (Network of Antimicrobial Resistance in *S. aureus* program), which was supported under the NIH-National Institute of Allergy and Infectious Diseases contract HHSN272200700055C. This work was supported in part through the computational and data resources and staff expertise provided by Scientific Computing and Data at the Icahn School of Medicine at Mount Sinai and supported by the Clinical and Translational Science Awards (CTSA) grant UL1TR004419 from the National Center for Advancing Translational Sciences. Research reported in this publication was also supported by the Office of Research Infrastructure of the National Institutes of Health under award number S10OD026880 and S10OD030463. The graphical abstract and Figure 1A were created using BioRender.com.

## Footnotes

Current affiliations: RT is currently in the Zuckerman Institute at Columbia University, New York, NY. II is currently at International Flavors & Fragrances, Palo Alto, CA. KAL is currently a Senior Scientist at Regeneron, Inc, Tarrytown, NY. RK is currently in the Dermatology department at Mount Sinai Icahn School of Medicine, New York, NY, USA. CZ is currently in the Department of Pathology and Microbiology at University of Nebraska Medical Center, Omaha, NE.

## References

1. Weber JT. Community-associated methicillin-resistant Staphylococcus aureus. Clin Infect Dis. 2005;41 Suppl 4:S269–72.

2. Herold BC, Immergluck LC, Maranan MC, Lauderdale DS, Gaskin RE, Boyle-Vavra S, et al. Community-acquired methicillin-resistant Staphylococcus aureus in children with no identified predisposing risk. JAMA. 1998;279(8):593–8.

3. Moran GJ, Krishnadasan A, Gorwitz RJ, Fosheim GE, McDougal LK, Carey RB, et al. Methicillin-resistant S. aureus infections among patients in the emergency department. N Engl J Med. 2006;355(7):666–74.

4. Li M, Du X, Villaruz AE, Diep BA, Wang D, Song Y, et al. MRSA epidemic linked to a quickly spreading colonization and virulence determinant. Nat Med. 2012;18(5):816–9.

5. Copin R, Sause WE, Fulmer Y, Balasubramanian D, Dyzenhaus S, Ahmed JM, et al. Sequential evolution of virulence and resistance during clonal spread of community-acquired methicillin-resistant Staphylococcus aureus. Proc Natl Acad Sci U S A. 2019.

6. Dini M, Shokoohizadeh L, Jalilian FA, Moradi A, and Arabestani MR. Genotyping and characterization of prophage patterns in clinical isolates of Staphylococcus aureus. BMC Res Notes. 2019;12(1):669.

7. McCarthy AJ, Witney AA, and Lindsay JA. Staphylococcus aureus temperate bacteriophage: carriage and horizontal gene transfer is lineage associated. Front Cell Infect Microbiol. 2012;2:6.

8. Sweet T, Jr., Sindi S, and Sistrom M. Going through phages: a computational approach to revealing the role of prophage in Staphylococcus aureus. Access Microbiol. 2023;5(6):acmi000424.

9. Xia G, and Wolz C. Phages of Staphylococcus aureus and their impact on host evolution. Infect Genet Evol. 2014;21:593–601.

10. Copin R, Shopsin B, and Torres VJ. After the deluge: mining Staphylococcus aureus genomic data for clinical associations and host-pathogen interactions. Curr Opin Microbiol. 2018;41:43–50.

11. Bae T, Baba T, Hiramatsu K, and Schneewind O. Prophages of Staphylococcus aureus Newman and their contribution to virulence. Mol Microbiol. 2006;62(4):1035–47.

12. Ingmer H, Gerlach D, and Wolz C. Temperate Phages of Staphylococcus aureus. Microbiol Spectr. 2019;7(5).

13. Kaneko J, Kimura T, Narita S, Tomita T, and Kamio Y. Complete nucleotide sequence and molecular characterization of the temperate staphylococcal bacteriophage phiPVL carrying Panton-Valentine leukocidin genes. Gene. 1998;215(1):57–67.

14. Gillet Y, Issartel B, Vanhems P, Fournet JC, Lina G, Bes M, et al. Association between Staphylococcus aureus strains carrying gene for Panton-Valentine leukocidin and highly lethal necrotising pneumonia in young immunocompetent patients. Lancet. 2002;359(9308):753-9.

15. Otter JA, Kearns AM, French GL, and Ellington MJ. Panton-Valentine leukocidin-encoding bacteriophage and gene sequence variation in community-associated methicillin-resistant Staphylococcus aureus. Clin Microbiol Infect. 2010;16(1):68–73.

16. Vandenesch F, Naimi T, Enright MC, Lina G, Nimmo GR, Heffernan H, et al. Community-acquired methicillin-resistant Staphylococcus aureus carrying Panton-Valentine leukocidin genes: worldwide emergence. Emerg Infect Dis. 2003;9(8):978–84.

17. Thurlow LR, Joshi GS, and Richardson AR. Virulence strategies of the dominant USA300 lineage of community-associated methicillin-resistant Staphylococcus aureus (CA-MRSA). FEMS Immunol Med Microbiol. 2012;65(1):5–22.

18. Balasubramanian D, Ohneck EA, Chapman J, Weiss A, Kim MK, Reyes-Robles T, et al. Staphylococcus aureus Coordinates Leukocidin Expression and Pathogenesis by Sensing Metabolic Fluxes via RpiRc. MBio. 2016;7(3).

19. Bonar EA, Bukowski M, Hydzik M, Jankowska U, Kedracka-Krok S, Groborz M, et al. Joint Genomic and Proteomic Analysis Identifies Meta-Trait Characteristics of Virulent and Non-virulent Staphylococcus aureus Strains. Front Cell Infect Microbiol. 2018;8:313.

20. Altman DR, Sullivan MJ, Chacko KI, Balasubramanian D, Pak TR, Sause WE, et al. Genome Plasticity of agr-Defective Staphylococcus aureus during Clinical Infection. Infect Immun. 2018;86(10).

21. Li M, Diep BA, Villaruz AE, Braughton KR, Jiang X, DeLeo FR, et al. Evolution of virulence in epidemic community-associated methicillin-resistant Staphylococcus aureus. Proc Natl Acad Sci U S A. 2009;106(14):5883–8.

22. Little JW. Phages. 2005:37-54.

23. Adhikari S, and Curtis PD. DNA methyltransferases and epigenetic regulation in bacteria. FEMS Microbiol Rev. 2016;40(5):575–91.

24. Blow MJ, Clark TA, Daum CG, Deutschbauer AM, Fomenkov A, Fries R, et al. The Epigenomic Landscape of Prokaryotes. PLoS Genet. 2016;12(2):e1005854.

25. Nye TM, Fernandez NL, and Simmons LA. A positive perspective on DNA methylation: regulatory functions of DNA methylation outside of host defense in Gram-positive bacteria. Crit Rev Biochem Mol Biol. 2020;55(6):576–91.

26. Fang G, Munera D, Friedman DI, Mandlik A, Chao MC, Banerjee O, et al. Genome-wide mapping of methylated adenine residues in pathogenic Escherichia coli using single-molecule real-time sequencing. Nat Biotechnol. 2012;30(12):1232–9.

27. Chen J, Yoong P, Ram G, Torres VJ, and Novick RP. Single-copy vectors for integration at the SaPI1 attachment site for Staphylococcus aureus. Plasmid. 2014;76:1–7.

28. Kossykh VG, Schlagman SL, and Hattman S. Conserved sequence motif DPPY in region IV of the phage T4 Dam DNA-[N-adenine]-methyltransferase is important for S-adenosyl-L-methionine binding. Nucleic Acids Res. 1993;21(15):3563–6.

29. Siwek W, Czapinska H, Bochtler M, Bujnicki JM, and Skowronek K. Crystal structure and mechanism of action of the N6-methyladenine-dependent type IIM restriction endonuclease R.DpnI. Nucleic Acids Res. 2012;40(15):7563–72.

30. Mader U, Nicolas P, Depke M, Pane-Farre J, Debarbouille M, van der Kooi-Pol MM, et al. Staphylococcus aureus Transcriptome Architecture: From Laboratory to Infection-Mimicking Conditions. PLoS Genet. 2016;12(4):e1005962.

31. Ahmed S, Meghji S, Williams RJ, Henderson B, Brock JH, and Nair SP. Staphylococcus aureus fibronectin binding proteins are essential for internalization by osteoblasts but do not account for differences in intracellular levels of bacteria. Infect Immun. 2001;69(5):2872–7.

32. Dziewanowska K, Patti JM, Deobald CF, Bayles KW, Trumble WR, and Bohach GA. Fibronectin binding protein and host cell tyrosine kinase are required for internalization of Staphylococcus aureus by epithelial cells. Infect Immun. 1999;67(9):4673–8.

33. Peacock SJ, Foster TJ, Cameron BJ, and Berendt AR. Bacterial fibronectin-binding proteins and endothelial cell surface fibronectin mediate adherence of Staphylococcus aureus to resting human endothelial cells. Microbiology (Reading). 1999;145 ( Pt 12):3477–86.

34. Fitzgerald JR, Loughman A, Keane F, Brennan M, Knobel M, Higgins J, et al. Fibronectin-binding proteins of Staphylococcus aureus mediate activation of human platelets via fibrinogen and fibronectin bridges to integrin GPIIb/IIIa and IgG binding to the FcgammaRIIa receptor. Mol Microbiol. 2006;59(1):212–30.

35. Herman-Bausier P, El-Kirat-Chatel S, Foster TJ, Geoghegan JA, and Dufrene YF. Staphylococcus aureus Fibronectin-Binding Protein A Mediates Cell-Cell Adhesion through Low-Affinity Homophilic Bonds. mBio. 2015;6(3):e00413–15.

36. Vergara-Irigaray M, Valle J, Merino N, Latasa C, Garcia B, Ruiz de Los Mozos I, et al. Relevant role of fibronectin-binding proteins in Staphylococcus aureus biofilm-associated foreign-body infections. Infect Immun. 2009;77(9):3978–91.

37. O’Neill E, Pozzi C, Houston P, Humphreys H, Robinson DA, Loughman A, et al. A novel Staphylococcus aureus biofilm phenotype mediated by the fibronectin-binding proteins, FnBPA and FnBPB. J Bacteriol. 2008;190(11):3835–50.

38. Que YA, Haefliger JA, Piroth L, Francois P, Widmer E, Entenza JM, et al. Fibrinogen and fibronectin binding cooperate for valve infection and invasion in Staphylococcus aureus experimental endocarditis. J Exp Med. 2005;201(10):1627–35.

39. Shinji H, Yosizawa Y, Tajima A, Iwase T, Sugimoto S, Seki K, et al. Role of fibronectin-binding proteins A and B in in vitro cellular infections and in vivo septic infections by Staphylococcus aureus. Infect Immun. 2011;79(6):2215–23.

40. Arciola CR, Campoccia D, and Montanaro L. Implant infections: adhesion, biofilm formation and immune evasion. Nat Rev Microbiol. 2018;16(7):397–409.

41. Kwiecinski J, Jin T, and Josefsson E. Surface proteins of Staphylococcus aureus play an important role in experimental skin infection. APMIS. 2014;122(12):1240–50.

42. Gries CM, Biddle T, Bose JL, Kielian T, and Lo DD. Staphylococcus aureus Fibronectin Binding Protein A Mediates Biofilm Development and Infection. Infect Immun. 2020;88(5).

43. Tuon FF, Suss PH, Telles JP, Dantas LR, Borges NH, and Ribeiro VST. Antimicrobial Treatment of Staphylococcus aureus Biofilms. Antibiotics (Basel). 2023;12(1).

44. Stoodley P, Nistico L, Johnson S, Lasko LA, Baratz M, Gahlot V, et al. Direct demonstration of viable Staphylococcus aureus biofilms in an infected total joint arthroplasty. A case report. J Bone Joint Surg Am. 2008;90(8):1751–8.

45. Elgharably H, Hussain ST, Shrestha NK, Blackstone EH, and Pettersson GB. Current Hypotheses in Cardiac Surgery: Biofilm in Infective Endocarditis. Semin Thorac Cardiovasc Surg. 2016;28(1):56–9.

46. Brady RA, Leid JG, Calhoun JH, Costerton JW, and Shirtliff ME. Osteomyelitis and the role of biofilms in chronic infection. FEMS Immunol Med Microbiol. 2008;52(1):13–22.

47. May JG, Shah P, Sachdeva L, Micale M, Kruper GJ, Sheyn A, et al. Potential role of biofilms in deep cervical abscess. Int J Pediatr Otorhinolaryngol. 2014;78(1):10–3.

48. Lacey KA, Serpas L, Makita S, Wang Y, Rashidfarrokhi A, Soni C, et al. Secreted mammalian DNases protect against systemic bacterial infection by digesting biofilms. J Exp Med. 2023;220(6).

49. Mansour SC, Pletzer D, de la Fuente-Nunez C, Kim P, Cheung GYC, Joo HS, et al. Bacterial Abscess Formation Is Controlled by the Stringent Stress Response and Can Be Targeted Therapeutically. EBioMedicine. 2016;12:219–26.

50. Sugimoto S, Sato F, Miyakawa R, Chiba A, Onodera S, Hori S, et al. Broad impact of extracellular DNA on biofilm formation by clinically isolated Methicillin-resistant and - sensitive strains of Staphylococcus aureus. Sci Rep. 2018;8(1):2254.

51. Malachowa N, and DeLeo FR. Mobile genetic elements of Staphylococcus aureus. Cell Mol Life Sci. 2010;67(18):3057–71.

52. Murphy J, Mahony J, Ainsworth S, Nauta A, and Sinderen Dv. Bacteriophage Orphan DNA Methyltransferases: Insights from Their Bacterial Origin, Function, and Occurrence. Applied and Environmental Microbiology. 2013;79(24):7547–55.

53. Chabelskaya S, Gaillot O, and Felden B. A Staphylococcus aureus small RNA is required for bacterial virulence and regulates the expression of an immune-evasion molecule. PLoS Pathog. 2010;6(6):e1000927.

54. Pichon C, and Felden B. Small RNA genes expressed from Staphylococcus aureus genomic and pathogenicity islands with specific expression among pathogenic strains. Proc Natl Acad Sci U S A. 2005;102(40):14249–54.

55. Chao MC, Zhu S, Kimura S, Davis BM, Schadt EE, Fang G, et al. A Cytosine Methyltransferase Modulates the Cell Envelope Stress Response in the Cholera Pathogen [corrected]. PLoS Genet. 2015;11(11):e1005666.

56. Bryant JM, Brown KP, Burbaud S, Everall I, Belardinelli JM, Rodriguez-Rincon D, et al. Stepwise pathogenic evolution of Mycobacterium abscessus. Science. 2021;372(6541).

57. Sadykov MR. Restriction-Modification Systems as a Barrier for Genetic Manipulation of Staphylococcus aureus. Methods Mol Biol. 2016;1373:9–23.

58. Pragman AA, and Schlievert PM. Virulence regulation in Staphylococcus aureus: the need for in vivo analysis of virulence factor regulation. FEMS Immunol Med Microbiol. 2004;42(2):147–54.

59. Goerke C, Koller J, and Wolz C. Ciprofloxacin and trimethoprim cause phage induction and virulence modulation in Staphylococcus aureus. Antimicrob Agents Chemother. 2006;50(1):171–7.

60. Krausz KL, and Bose JL. Bacteriophage Transduction in Staphylococcus aureus: Broth-Based Method. Methods Mol Biol. 2016;1373:63–8.

61. Monk IR, Shah IM, Xu M, Tan MW, and Foster TJ. Transforming the untransformable: application of direct transformation to manipulate genetically Staphylococcus aureus and Staphylococcus epidermidis. MBio. 2012;3(2).

62. Thornton JA. Splicing by Overlap Extension PCR to Obtain Hybrid DNA Products. Methods Mol Biol. 2016;1373:43–9.

63. Fey PD, Endres JL, Yajjala VK, Widhelm TJ, Boissy RJ, Bose JL, et al. A genetic resource for rapid and comprehensive phenotype screening of nonessential Staphylococcus aureus genes. mBio. 2013;4(1):e00537–12.

64. Li H, and Durbin R. Fast and accurate short read alignment with Burrows-Wheeler transform. Bioinformatics. 2009;25(14):1754–60.

65. Quinlan AR, and Hall IM. BEDTools: a flexible suite of utilities for comparing genomic features. Bioinformatics. 2010;26(6):841–2.

66. Wickham H. ggplot2: Elegant Graphics for Data Analysis. https://ggplot2.tidyverse.org. 2023.

67. Zheng X, Marsman G, Lacey KA, Chapman JR, Goosmann C, Ueberheide BM, et al. The cell envelope of Staphylococcus aureus selectively controls the sorting of virulence factors. Nat Commun. 2021;12(1):6193.

68. Klopfenstein N, Cassat JE, Monteith A, Miller A, Drury S, Skaar E, et al. Murine Models for Staphylococcal Infection. Curr Protoc. 2021;1(3):e52.

69. Schneider CA, Rasband WS, and Eliceiri KW. NIH Image to ImageJ: 25 years of image analysis. Nat Methods. 2012;9(7):671–5.

70. Dyzenhaus S, Sullivan MJ, Alburquerque B, Boff D, van de Guchte A, Chung M, et al. MRSA lineage USA300 isolated from bloodstream infections exhibit altered virulence regulation. Cell Host Microbe. 2023;31(2):228–42 e8.

71. Tatusova T, DiCuccio M, Badretdin A, Chetvernin V, Nawrocki EP, Zaslavsky L, et al. NCBI prokaryotic genome annotation pipeline. Nucleic Acids Res. 2016;44(14):6614–24.

72. Livak KJ, and Schmittgen TD. Analysis of relative gene expression data using real-time quantitative PCR and the 2(-Delta Delta C(T)) Method. Methods. 2001;25(4):402–8.

73. Rona G, Miwatani-Minter B, Zhang Q, Goldberg HV, Kerzhnerman MA, Howard JB, et al. D-type cyclins regulate DNA mismatch repair in the G1 and S phases of the cell cycle, maintaining genome stability. bioRxiv. 2024.

74. Boles BR, Thoendel M, Roth AJ, and Horswill AR. Identification of genes involved in polysaccharide-independent Staphylococcus aureus biofilm formation. PLoS One. 2010;5(4):e10146.

75. Monk IR, Tree JJ, Howden BP, Stinear TP, and Foster TJ. Complete Bypass of Restriction Systems for Major Staphylococcus aureus Lineages. mBio. 2015;6(3):e00308–15.

76. Novick RP. Genetic systems in staphylococci. Methods Enzymol. 1991;204:587–636.

77. Benson MA, Lilo S, Wasserman GA, Thoendel M, Smith A, Horswill AR, et al. Staphylococcus aureus regulates the expression and production of the staphylococcal superantigen-like secreted proteins in a Rot-dependent manner. Mol Microbiol. 2011;81(3):659–75.

78. Geisinger E, George EA, Chen J, Muir TW, and Novick RP. Identification of ligand specificity determinants in AgrC, the Staphylococcus aureus quorum-sensing receptor. J Biol Chem. 2008;283(14):8930–8.

79. Anderson EE, Dyzenhaus S, Ilmain JK, Sullivan MJ, van Bakel H, and Torres VJ. SarS Is a Repressor of Staphylococcus aureus Bicomponent Pore-Forming Leukocidins. Infect Immun. 2023;91(4):e0053222.

