## Supplemental tables, figures, and methods for "Prophage-encoded methyltransferase drives adaptation of community-acquired methicillin-resistant *Staphylococcus aureus*"

**Supplemental Figures and Tables**

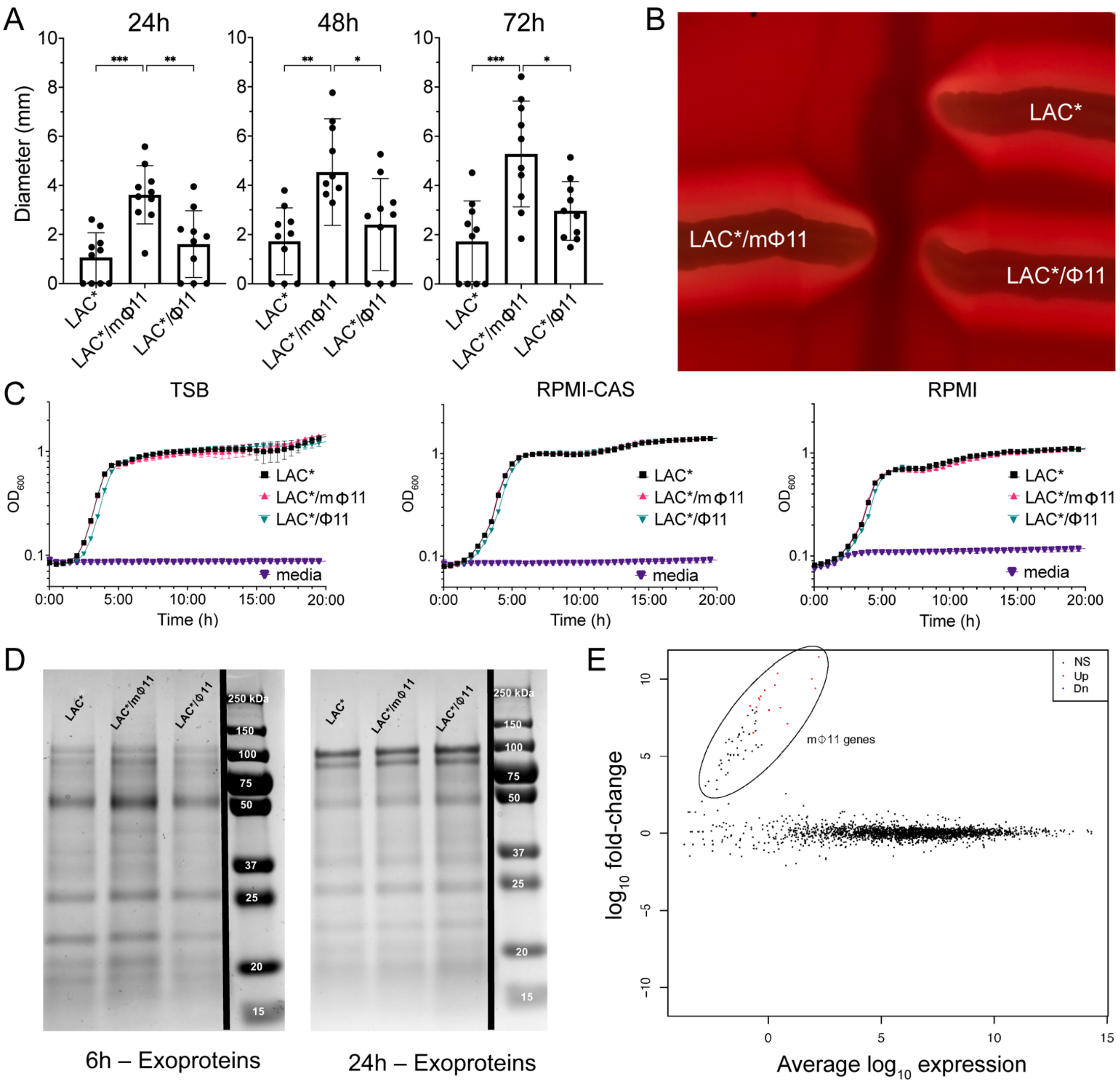


**Figure S1. Prophage mΦ11 increases skin abscess size but does not affect in vitro growth, exoprotein production, hemolysis, or transcription of non-phage genes. (A)** Skin abscess size. Skin abscess diameter of LAC* (N=10 abscesses), LAC*/mΦ11 (N=10

abscesses, strain BS989), and LAC*/Φ11 (N=10 abscesses, strain BS990) at the indicated time points after injection with ~$10^7$ CFU of bacteria per abscess. Data are results of one experiment that replicates prior findings (1). Data represent mean ± SD. Statistical significance was determined with the Kruskall-Wallis test and Dunn's multiple comparisons test, *$P \leq .05$, **$P \leq .01$, ***$P \leq .001$. **(B)** Hemolysis patterns. Cross-streaking of the indicated strains alongside the β-hemolysin producing strain RN4220 (streaked vertically) on a sheep blood agar plate; differentiation of the various hemolytic activities in *S. aureus* can be scored on sheep blood agar by virtue of their synergism with β-hemolysin. Hemolysis patterns were identical, irrespective of the presence or absence of the mosaic phage. **(C)** Growth curves of the indicated strains in various media. Growth was monitored by measuring $OD_{600}$. Data are the mean and SD of three biological replicates. Results are representative of two independent experiments. **(D)** Exoprotein profiles. Exoproteins profiles prepared from the culture supernatants of the indicated strains, analyzed by SDS-PAGE and staining with Coomassie Blue after growth for 6 h and 24 h. Black line indicates the lanes were non-contiguous with the ladder. Images represent two independent experiments. **(E)** Prophage mΦ11-mediated transcriptional changes. Scatter plot comparing log RNA transcript levels at late exponential (6 h) growth in TSB of LAC* and LAC*/mΦ11 strains. Orange data points represent genes with increased expression in LAC*/mΦ11 that met statistical significance ($P \leq .05$), and the circled area represents mΦ11 genes. Data shown is representative of two separate experiments.

Collectively, the data show that in vitro analyses do not correlate with abscess size or phage content, suggesting that the changes that accompany mΦ11-mediated enhanced virulence in the skin require host tissue-specific signals in vivo.

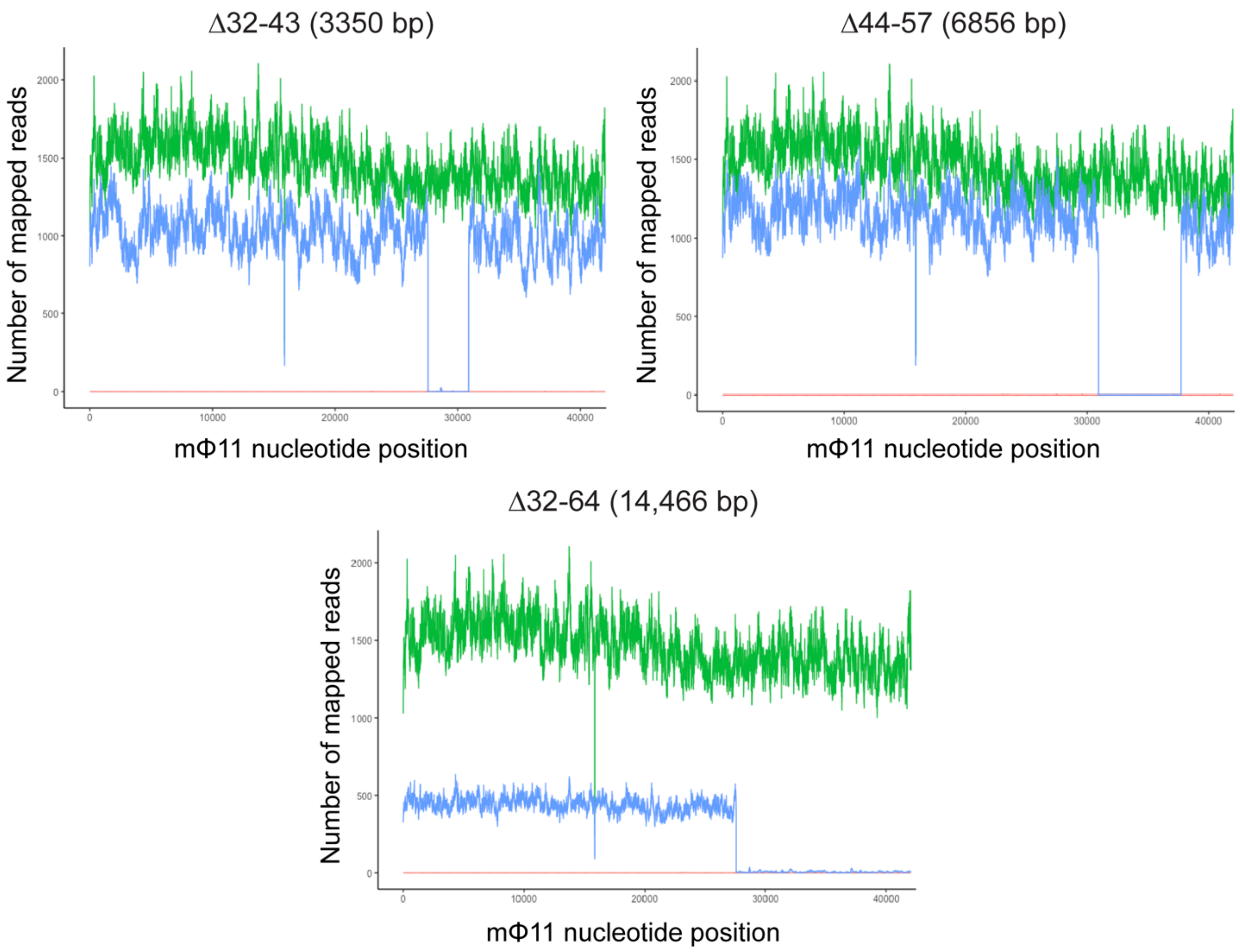


**Figure S2. Validation of en bloc deletion clones by whole genome sequencing.** Genomic comparison of LAC*/mΦ11 (green, strain BS989), LAC*/mΦ11 containing the indicated en bloc deletion (blue, strains RU42, RU47 and RU108), and control LAC* (red) strains using bedtools v2.30.0 (2) with the mΦ11 sequence as a reference. The sharp cutoffs to zero reads in the en bloc deletion strains correspond to the expected deletion locations in each strain, confirming the deletions.

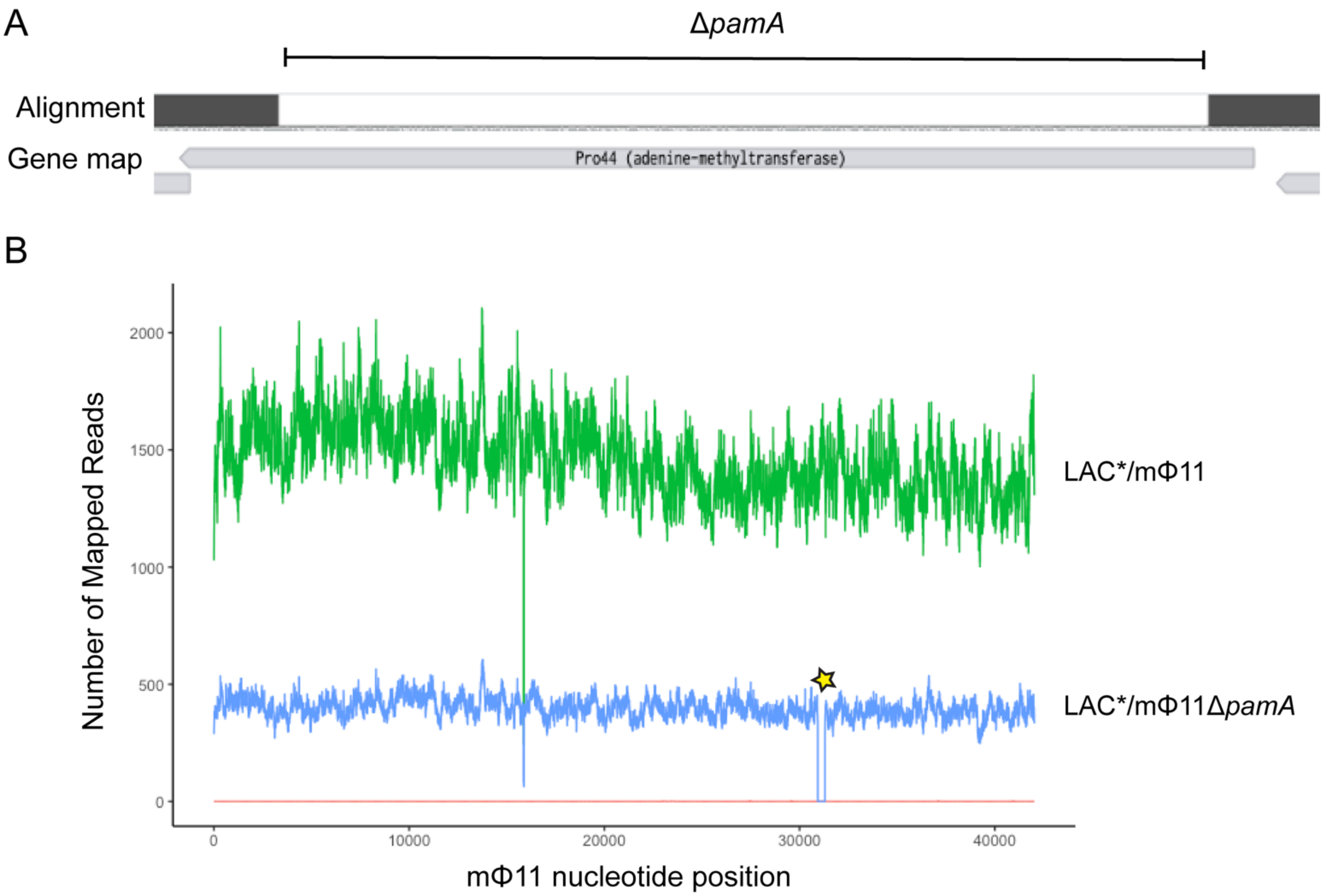


**Figure S3. Validation of unmarked, in-frame deletion of *pamA* by Sanger and whole genome sequencing. (A)** Alignment of *pamA* sequence from LAC*/mΦ11Δ*pamA* (strain RU39) obtained using Sanger Sequencing to reference mΦ11 using Benchling alignment tool (https://www.benchling.com/). Area highlighted by bracket, with no alignment to the reference gene map, indicates the 369 bp deletion in Δ*pamA*. **(B)** Whole genome sequencing comparison of LAC*/mΦ11 (green, strain BS989), USA300-LAC*/mΦ11Δ*pamA* (blue, strain RU39), and control LAC* (red) to reference mΦ11, performed as in Figure S2. Genomic comparison showing that all reads from LAC*/mϕ11 and LAC* control is present in LAC*/mΦ11Δ*pamA* except for gaps in coverage that correspond to the location of *pamA*, confirming the *pamA* deletion (highlighted by yellow star) and the absence of adventitious secondary mutations.

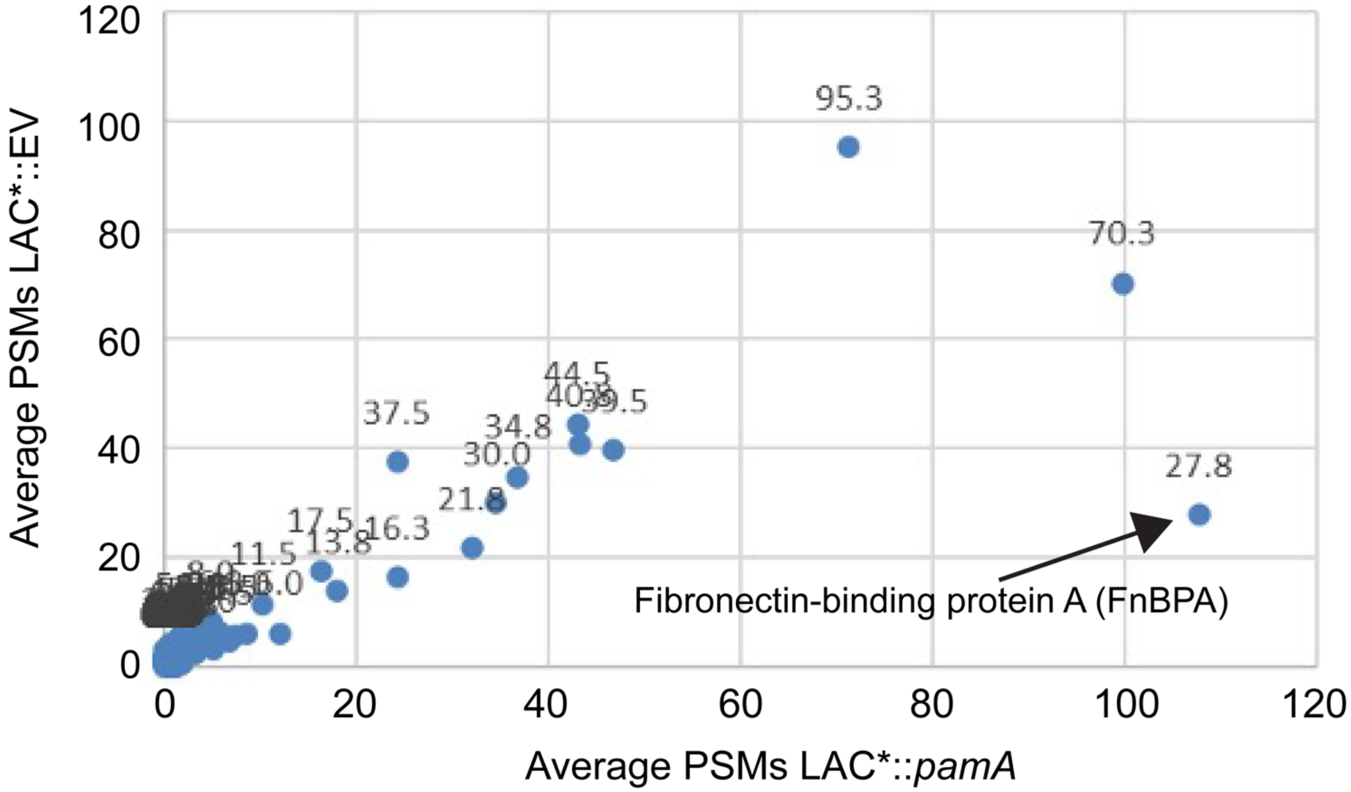


**Figure S4. Mass spectrometry suggests that FnBPA is the high molecular weight cell-wall-associated protein in LAC*::*pamA* biofilms.** The high molecular weight band in LAC*::*pamA* lanes (see **Figure 6B**) and corresponding high molecular weight areas of LAC*::EV lanes were extracted and analyzed by mass spectrometry. The average Peptide Spectral Matches (PSM) of LAC*::*pamA* (N=4 biological replicates, strain RU121) and LAC*::EV (N=4 biological replicates, strain RU129) are shown on the scatter plot. Data are pooled from two independent replicate experiments. The numbers above the data points correspond to the average PSMs in LAC*::EV. The data point representing FnBPA is highlighted with a black arrow.

| Start | End | Annotated product | Gene name | Motifs |
|---|---|---|---|---|
| 2056937 | 2057362 | adenine-methyltransferase | ph11Mos_00044 | Dam |
| 2057606 | 2057767 | Hypothetical protein | ph11Mos_00046 | no domain |
| 2057761 | 2058540 | DNA replication protein DnaC | ph11Mos_00047 | IstB_IS21 |
| 2058550 | 2059272 | Helix-turn-helix protein | ph11Mos_00048 | HTH_36 |
| 2059265 | 2059672 | HNH endonuclease | ph11Mos_00049 | HNH |
| 2059727 | 2060491 | AP2 domain protein | ph11Mos_00050 | AP2 |
| 2062530 | 2062751 | Hypothetical protein | ph11Mos_00054 | DUF2483 |
| 2063564 | 2063794 | Hypothetical protein | ph11Mos_00057 | no domain |

**Table S1. Candidate gene(s) for increased virulence localized by en bloc deletion experiments.** Table adapted from (1) with start and stop positions, predicted annotations, gene name and predicted protein motifs of the mΦ11 ORFs that did not share homology with Φ11 within the candidate “virulence block” (Δ44-57) identified by en bloc deletion experiments (see **Figure 1**).

### Supplemental Methods

#### A. Construction of bacterial strains

*Construction of en bloc and pamA deletion mutants.* In-frame, unmarked, deletions were engineered by amplifying 600-1000 base pair fragments flanking the target gene(s) using genomic DNA (gDNA) template from strain LAC*/mΦ11 (BS989) and the following oligonucleotide pairs [oRU1/oRU2 (upstream) and oRU3/oRU4 (downstream) to construct pRU1 for LAC*/mΦ11Δ*pamA* (RU39); oRU5/oRU6 (upstream) and oRU7/oRU8 (downstream) to construct pRU2 for LAC*/mΦ11Δ32-64 (RU42); oRU5/oRU6 (upstream) and oRU9/oRU4 (downstream) to construct pRU3 for LAC*/mΦ11Δ32-43 (RU47); oRU17/oRU2 (upstream) and oRU10/oRU11 (downstream) to construct pRU4 for LAC*/mΦ11Δ44-57 (RU108)] **(Table 1)**. Amplified fragments were purified with PCR clean-up kit (Qiagen, #28104) and were inserted into an inverse PCR amplified (oRU15/oRU16) pIMAY cloning plasmid (3) using Gibson assembly (New England Biolabs, #E2611S). Next, 5 µL of Gibson product was added to 50µL of chemically competent *E. coli* IM08, incubated 30 min on ice, heat shocked in 42°C water for 45s, then 950 µL LB was added for recovery incubation at 37°C with shaking at 250 rpm. After recovery, 100 µL of sample was plated on prewarmed LB plates with Cm and incubated at 37°C. Plasmids pRU1, pRU2, pRU3 and pRU4 were isolated from colonies using QIAprep Spin Miniprep kit (Qiagen, #27106). The cloning inserts were validated by Sanger sequencing (Psomagen, Inc.) of PCR amplification product using primers IM151/IM152 and the plasmids were electroporated into electrocompetent strain LAC*/mΦ11 (BS989) for allelic exchange using established protocols (3).

*Chromosomal insertion of constitutively expressed pamA.* For single-copy chromosomal insertion, a fragment containing the constitutive *sarA* promoter and *sod* ribosome binding site

($P_{sarA}$-*sod*RBS) was PCR amplified using DNA template from pOS1-$P_{SarA}$-*sod*RBS-GFP (4) and oligonucleotide pair oRU43/oRU44, and a fragment containing the full-length *pamA* gene was PCR amplified using gDNA template from strain BS989 and oligonucleotide pair oRU45/oRU46. The fragments were separated on 1% agarose gel and isolated and purified with QIAquick gel extraction kit (Qiagen, #28704) per manufacturers protocol. pJC1111, a suicide plasmid with Amp and Cd resistance markers that integrates at the *S. aureus* pathogenicity island 1 (SapI1) *att* site (5), was PCR amplified using oligonucleotide pair oRU41/oRU42 and the $P_{sarA}$-*sod*RBS and *pamA* amplification products were ligated into pJC1111 using Gibson assembly, generating pRU7. Transformants were selected on LB/Amp plates and validated by sequencing using primers oRU80/oRU81. Once the insert was confirmed, pRU7 was isolated using a QIAprep Spin Miniprep kit (Qiagen, #27106) and transformed into strain BS656 for chromosomal integration using published protocols (5), generating strain RU116. To generate control strains, pJC1111 was transformed into strain BS656 for chromosomal integration, generating strain RU120. Chromosomal integrants at the SapI1 chromosomal position were selected with $CdCl_2$. Phage 80α lysates of RU116 were used to transduce LAC* and RU39, generating RU121 and RU131, respectively. Phage 80α lysates of RU120 were used to transduce LAC*, RU39, and BS989 generating empty vector (EV) control strains RU129, RU128, and RU138 respectively. The $P_{sarA}$-*sod*RBS-*pamA* and EV inserts were validated by Sanger sequencing using primers oRU80/oRU81.

*Chromosomal insertion of constitutively expressed pamA point mutants.* Point mutants were created using complementary oligonucleotides engineered to contain the desired *pamA* point mutation [oRU70/oRU43 (upstream) and oRU71/oRU46 (downstream) to construct pRU9 for *pamA*P65A strain RU161; oRU72/oRU43 (upstream) and oRU73/oRU46 (downstream) to construct pRU10 for *pamA*P65T mutant strain RU162; oRU74/oRU43 (upstream) and oRU75/oRU46 (downstream) to construct pRU11 for PamAP66A mutant (strain RU164)] to amplify fragments from pJC1111::$P_{sarA}$-*sod*RBS-*pamA* template DNA. The fragments were gel purified using QIAquick gel extraction kit (Qiagen, #28704) and joined with overlap extension (OE) PCR (6) using oligonucleotide pair oRU43/oRU46. After OE-PCR, the fragments were inserted into pJC1111 using Gibson assembly, transformed into competent *E. Coli* DH5α (New England Biolabs, #C2987H) per manufacturer instructions and plated on LB/Amp plates. Colony PCR (primers oRU80/oRU81) of colonies growing on LB/Amp plates confirmed the insert and the product was sent for Sanger sequencing. Once the desired point-mutated insert was confirmed, pRU9, pRU10, and pRU11 were harvested from *E. Coli* DH5α using Qiagen miniprep kit, electroporated into competent *S. aureus* strain BS656 (resulting in strains RU152, RU154, and RU156, respectively), and transduced into strain LAC* using phage 80α (resulting in strains RU161, RU162, and RU164, respectively). Transductants were selected on TSA/Cd plates, underwent colony PCR using oligonucleotide pair oRU80/oRU81 and Sanger sequencing of the amplified product confirmed the desired point mutation. The inactivation of *pamA* was confirmed with DpnI digestion as described below.

*Fibronectin-binding protein A (fnbA) transposon mutants*. Phage 80α lysate of Nebraska Transposon Mutant Library (7) strain NE186 (*fnbA*::*bursa,* Erm) was used to transduce RU121, RU129, and BS989, to generate RU169, RU170, and RU171, respectively. PCR amplification using primers to amplify the *fnbA* transposon insertion site (oRU84/oRU85) confirmed the insertion.

**B. Protein identification by mass spectrometry**

After digestion, the solution containing the peptides was transferred into a new Eppendorf tube and the gels washed via shaking with a 1:2 (vol/vol) 5% formic acid/ acetonitrile extraction buffer for 15 min at 37°C. The extraction buffer was removed and combined with the previous aspirated solution and the process repeated two more times. The combined solutions were dried using a SpeedVac (Thermo Savant). The dried sample was reconstituted in 0.1% TFA and desalted using C18 microspin columns (Harvard apparatus (Millipore)). The bound peptides were rinsed three times with 0.1% TFA and one time with 0.5% acetic acid and elute off the microspin column using 40 μL 40% acetonitrile in 0.5% acetic acid followed by the addition of 40 μL 80% acetonitrile in 0.5% acetic acid. The organic solvent was removed using a SpeedVac concentrator and the sample was reconstituted in 0.5% acetic acid. For mass spectrometry (MS) analysis, an aliquot of each sample (1/10) was loaded onto an Acclaim PepMap trap column (2 cm x 75 μm) in line with an EASY-Spray analytical column (50 cm x 75 μm ID PepMap C18, 2 μm bead size) using the autosampler of an EASY-nLC 1200 HPLC (Thermo Scientific). Solvent A was 2% acetonitrile in 0.5% acetic acid and solvent B was 80% acetonitrile in 0.5% acetic acid. The sample was gradient eluted in the Orbitrap Eclipse (Thermo Fisher). The gradient was held for 5 min at 5% solvent B, ramped in 60 min to 35% solvent B, in 10 min to 45% solvent B, and in another 10 min to 100% solvent B. High resolution full MS spectra were acquired with a resolution of 120,000, an AGC target of 4e5, with a maximum ion time of 50 ms, and scan range of 400 to 1500 m/z. Following each full MS scan, precursor ions with charge states between 2-5 were considered for fragmentation for a cycle duration of 3 seconds. MS/MS HCD spectra were collected with a resolution of 30,000, an AGC target of 2e5, maximum injection time of 200 ms, one microscan, 2 m/z isolation window, fixed first mass of 150, and normalized collision energy (NCE) of 27. Dynamic exclusion was set to 30 seconds. The MS/MS spectra were searched against a *S. aureus* USA300 database and common contaminants (cRAP) using Sequest within Proteome Discoverer 1.4. using the following settings: oxidized methionine (M), and deamidation (asparagine and glutamine) were selected as variable modifications, and carbamidomethyl (C) as fixed modifications; precursor and fragment mass tolerance was set to 10 ppm. The data was filtered to better than 1% peptide and protein FDR searched against a decoy database. Only proteins with at least two different peptides were considered for downstream analysis.

### References


1. Copin R, Sause WE, Fulmer Y, Balasubramanian D, Dyzenhaus S, Ahmed JM, et al. Sequential evolution of virulence and resistance during clonal spread of community-acquired methicillin-resistant Staphylococcus aureus. *Proc Natl Acad Sci U S A.* 2019.
2. Quinlan AR, and Hall IM. BEDTools: a flexible suite of utilities for comparing genomic features. *Bioinformatics.* 2010;26(6):841-2.
3. Monk IR, Shah IM, Xu M, Tan MW, and Foster TJ. Transforming the untransformable: application of direct transformation to manipulate genetically Staphylococcus aureus and Staphylococcus epidermidis. *MBio.* 2012;3(2).
4. Benson MA, Lilo S, Wasserman GA, Thoendel M, Smith A, Horswill AR, et al. Staphylococcus aureus regulates the expression and production of the staphylococcal superantigen-like secreted proteins in a Rot-dependent manner. *Mol Microbiol.* 2011;81(3):659-75.
5. Chen J, Yoong P, Ram G, Torres VJ, and Novick RP. Single-copy vectors for integration at the SaPI1 attachment site for Staphylococcus aureus. *Plasmid.* 2014;76:1-7.
6. Thornton JA. Splicing by Overlap Extension PCR to Obtain Hybrid DNA Products. *Methods Mol Biol.* 2016;1373:43-9.

7. Fey PD, Endres JL, Yajjala VK, Widhelm TJ, Boissy RJ, Bose JL, et al. A genetic resource for rapid and comprehensive phenotype screening of nonessential Staphylococcus aureus genes. *mBio.* 2013;4(1):e00537-12.
